# Large-scale Integrative Taxonomy (LIT): resolving the data conundrum for dark taxa

**DOI:** 10.1101/2021.04.13.439467

**Authors:** Emily Hartop, Amrita Srivathsan, Fredrik Ronquist, Rudolf Meier

**Affiliations:** Zoology Department, Stockholm University, Stockholm, Sweden; Station Linné, Öland, Sweden; Department of Biological Sciences, National University of Singapore, Singapore; Department of Bioinformatics and Genetics, Swedish Museum of Natural History, Stockholm, Sweden

## Abstract

New, rapid, accurate, scalable, and cost-effective species discovery and delimitation methods are needed for tackling “dark taxa”, that we here define as clades for which <10% of all species are described and the estimated diversity exceeds 1000 species. Species delimitation for these taxa should be based on multiple data sources (“integrative taxonomy”) but collecting multiple types of data risks impeding a discovery process that is already too slow. We here develop explicit methods to avoid this by applying Large-scale Integrative Taxonomy (LIT). Preliminary species hypotheses are generated based on inexpensive data that are obtained quickly and cost-effectively in a technical exercise. The validation step is then based on a more expensive type of data that are only obtained for specimens selected based on objective criteria. We here use this approach to sort 18 000 scuttle flies (Diptera: Phoridae) from Sweden into 315 preliminary species hypotheses based on NGS barcode (313bp) clusters. These clusters were subsequently tested with morphology and used to develop quantitative indicators for predicting which barcode clusters are in conflict with morphospecies. For this purpose, we first randomly selected 100 clusters for in-depth validation with morphology. Afterwards, we used a linear model to demonstrate that the best predictors for conflict between barcode clusters and morphology are maximum p-distance within the cluster and cluster stability across different clustering thresholds. A test of these indicators using the 215 remaining clusters reveals that these predictors correctly identify all clusters that conflict with morphology. The morphological validation step in our study involved only 1 039 specimens (5.8% of all specimens), but a newly proposed simplified protocol would only require the study of 915 (5.1%: 2.5 specimens per species), as we show that clusters without signatures of incongruence can be validated by only studying two specimens representing the most divergent haplotypes. To test the generality of our results across different barcode clustering techniques, we establish that the levels of conflict are similar across Objective Clustering (OC), Automatic Barcode Gap Discovery (ABGD), Poisson Tree Processes (PTP) and Refined Single Linkage (RESL) (used by Barcode of Life Data System (BOLD) to assign Barcode Index Numbers (BINs)). OC and ABGD achieved a maximum congruence score with morphology of 89% while PTP was slightly less effective (84%). RESL could only be tested for a subset of the specimens because the algorithm is not public. BINs based on 277 of the original 1 714 haplotypes were 86% congruent with morphology while the values were 89% for OC, 74% for PTP, and 72% for ABGD.

## Introduction

> *“I saw with regret, (and all scientific men have shared this feeling) that whilst the number of accurate instruments was daily increasing, we were still ignorant”*
>
> — *— Alexander von Humboldt*

In a recent report, global reinsurance giant Swiss Re concluded that biodiversity and ecosystem services (BES) “underpin all economic activity in our societies globally…55% of global GDP is moderately or highly dependent on BES” (Swiss Re Institute 2020). At the same time, the world of ESG (Environmental, Social and Governance) investing has hit mainstream with biodiversity as a key topic (Kishan and Marsh 2021). Natural capital is reliant on biodiversity and ecosystem services that are, in turn, critically dependent on functionally diverse invertebrate groups like insects. They contribute a wide range of ecosystem services (Losey and Vaughan 2006), comprise over half of described species (Chapman 2009), and are hosts for millions of unique bacterial species, nematodes, and mites (Larsen et al. 2017). The economic value of insects in the United States alone is estimated to exceed 57 billion USD annually (Losey and Vaughan 2006). Insects may comprise most of the described species on earth and be critical to the functioning of our planet, but the described diversity constitutes only a small fraction of the true diversity. This highlights the need for completing one of the great incomplete tasks in science, an inventory of all of life. This mission requires the development of efficient tools for delimiting and identifying invertebrates, given that—despite centuries of study—most eukaryotic species remain unknown to science (Mora et al. 2011). We lack even a consensus estimate of the number of species on our planet, yet alone a clear picture of their abundances, composition, ranges, functions, and forms (Mora et al. 2011; Locey and Lennon 2016).

The bulk of the planet’s unknown diversity is in neglected groups, some of which are so diverse that a reasonably precise estimate of true species numbers is currently impossible. Such clades used to be called “open-ended taxa” (Bickel 2009), but recently the term “dark taxa” is more commonly used, although it was originally coined for the growing number of sequences in GenBank that were not linked to formal scientific names (Page 2011, 2016). As used today, “dark taxa” refers to species-rich taxa of small body size for whom most of the species-level diversity is undescribed (Hausmann et al. 2020). Here, we accept the current usage, but also propose that it should only be applied to taxa for which the undescribed fauna is estimated to exceed the described fauna by at least one order of magnitude and the total diversity exceeds 1000 species.

Dark taxa are so abundant that they should be included in any holistic biodiversity assessment. For example, we estimate that on average each of the 36 two-week Malaise trap samples from Sweden studied here included 1000 phorids. If 10 traps had been run for two weeks in each of the 450 000 km^2^ of Sweden, 4.5 billion specimens would have obtained which surely would have only been well below 1% of the living flies in the habitats given that the traps are only 1.1 m wide. This means that at any point in time, there must be well over a trillion phorids active in Sweden. They represent substantial biomass and have numerous albeit largely unknown roles in the country’s ecosystems. Any number of biological services may rely on phorids. This includes those species that live in the high-latitude regions that are already witnessing visible effects of climate change; yet we know nothing about the natural history of these species. Contrast this with what we know about a single charismatic insect species like the Large Blue butterfly (*Phengaris arion*), also resident in Sweden. Its distribution, life history, associations, and behaviours are all well documented, and this single species has been the subject of sustained conservation and reintroduction campaigns across multiple countries (Thomas 1995, Andersen et al. 2014). It is unlikely that the Large Blue is so critical to ecosystems that they will collapse without it, but the Large Blue is an important symbol of conservation biology, a charismatic taxon that resonates with us (Yong 2009). We are not arguing against the attention paid to such species, but we here emphasize is that we need a new field in zoology that focuses on the taxonomy, systematics, and ecology of dark taxa. The reason has been succinctly summarised by Curtis for microbes where he referenced another large blue: “For if the last blue whale choked to death on the last panda, it would be disastrous but not the end of the world. But if we accidentally poisoned the last two species of ammonia-oxidizers, that would be another matter. It could be happening now and we wouldn’t even know...” (Curtis 2006).

Tackling these groups with traditional taxonomic techniques has been very slow because a single site can yield thousands of specimens belonging to hundreds of species (Puillandre et al. 2012; Srivathsan et al. 2019). Usually, most specimens belong to a handful of common species, making the discovery and identification of new or rare species much like finding a needle in the proverbial haystack. In other samples, the situation is more like finding a needle among many other needles, with high species numbers at more even abundances (Brown 2021). Examining samples using traditional techniques requires a great deal of time, expertise, and a particular skillset (slide mounting, dissections, etc). Additionally, the sheer quantity of species makes the task of recalling details of morphospecies difficult. Who can remember what species number 34 looked like by the time you get to morphospecies number 459? This is challenging even with voluminous notes, detailed drawings, and repeated referrals to voucher specimens. The result of these challenges has been “cherrypicking”, where a few morphologically distinct and/or species-poor lineages within hyperdiverse taxa are targeted for description while the most diverse subclades remain unstudied. Unfortunately, this practice is of limited value for advancing the knowledge of dark taxa in the era of biodiversity decline.

The proposal of DNA barcoding for the identification of organisms (Hebert et al. 2003) and DNA taxonomy (Tautz et al. 2003; Blaxter 2004; Vogler and Monaghan 2007) in the early 2000s highlighted the potential of DNA sequences for accelerating species discovery and species identification. However, the widespread use of DNA barcodes remained too expensive for large-scale implementations until Sanger sequencing was replaced with various 2^nd^ and 3^rd^ generation sequencing technologies. Only recently have next generation sequencing barcodes (NGS barcodes) become sufficiently cheap and easy to obtain for truly reversing the traditional workflow of first sorting specimens with morphology and then collecting DNA sequences for a select subset of specimens later (Puillandre et al. 2012; Kekkonen and Hebert 2014; Wang et al. 2018; Yeo et al. 2020). The analysis of such large barcode datasets has revealed that 10-20% of all barcode clusters differ depending on the method and parameters used for molecular species delimitation (Yeo et al. 2020), and in some taxa incongruence can be much higher (Kekkonen and Hebert 2014). This is not surprising because a single short barcode marker contains a limited amount of information relevant to species boundaries (Kwong et al. 2012; Pentinsaari et al. 2016). This means that data for multiple character systems are needed for delimiting species (Integrative taxonomy: Dayrat 2005; Padial and Miralles 2010; Schlick-Steiner et al. 2010; Puillandre et al. 2012; Ratnasingham and Hebert 2013; Zhang et al. 2013; Pante et al. 2015; Vitecek et al. 2017).

Despite this recognised need, efficient quantitative approaches to integrative taxonomy are still underdeveloped. Integrative taxonomy has mostly used traditional morphospecies sorting followed by barcoding of a few representative specimens and then focused more on description than on efficient delimitation (Butcher et al. 2012; Riedel et al. 2013; Lücking et al. 2016). Alternatively, some authors have started to describe species based on COI clusters alone. However, this is known to yield incorrect boundaries for a significant number of species and this practice has thus been rejected by researchers who otherwise embrace molecular data for taxonomic purposes (Puillandre et al. 2012; Ratnasingham and Hebert 2013; Zhang et al. 2013; Srivathsan et al. 2021).

Large-scale integrative taxonomy involving DNA barcodes was first proposed by Puillandre et al. (2012) who obtained *COI* barcodes from 1000 specimens and derived primary species hypotheses using multiple molecular delimitation methods (Automated Barcode Gap Discovery and General Mixed Yule Coalescence Method). The cluster conflict between methods was visualised, and other data sources (nuclear, morphological, and geographic) were used to test the primary hypotheses. However, this raises the issue of what we call the “data conundrum” for the integrative taxonomy of dark taxa. Collecting multiple types of data for all specimens yields high quality species limits, but also dramatically slows down taxonomic progress, while acceleration is what is needed. Puillandre et al. (2012) used congruence between barcode clusters obtained with different molecular delimitation methods as a first indication of whether a primary species hypothesis was likely to be valid. However, this approach has the downside that misleading signal in the molecular data would not be detected because all molecular species delimitation methods would be likely to yield congruent barcode clusters. We here pursue a different approach. We demonstrate that one can use specific properties of barcode clusters (e.g., maximum intra-cluster p-distance) to predict conflict with morphology. The ability to predict conflict is then used to develop an explicit specimen subsampling scheme that allows for picking only a moderately small number of specimens for which the second type of data must be collected in order to test weak primary species hypotheses.

Here, we propose Large-scale Integrative Taxonomy (LIT), a systematic approach to rapid species delimitation based on the reverse workflow and designed to handle dark taxa. The core goal of LIT is the use of multiple data sources for delimiting species without slowing down the overall species discovery and delimitation processes. This can be achieved by first generating preliminary species-level hypotheses based on data that can be acquired quickly and in semi-automated ways. Currently, the best choice is NGS barcodes because of the development of cost-effective high-throughput individual-specimen sequencing techniques (Hebert et al. 2017; Srivathsan et al. 2018, 2019; Wang et al. 2018). However, it appears likely that in the future other types of data (e.g., high throughput imaging) could replace or complement NGS barcoding. The species-level hypotheses derived from these primary data are then used to physically sort specimens and guide the selection of specimens for followup study using a second type of data to validate the preliminary species hypotheses (Puillandre et al. 2012). The second type of data can be more expensive and/or require more highly skilled manpower because they only need to be acquired for a small subset of specimens. Typical examples of such data would be morphology or DNA sequences from nuclear markers. Critically, LIT identifies clusters generated with the primary data source that are most likely to be problematic, allowing efforts to be focused where needed.

Here, we illustrate how LIT can be used for solving a taxonomic problem that was too large for the application of traditional techniques. This would have meant that these samples would have been ignored or only morphologically unusual species would have been “cherry-picked” for study. We use LIT to discover the species-level diversity in 18 000 specimens of scuttle flies (Diptera: Phoridae) from Sweden by first generating NGS barcodes and then creating barcode clusters. We then study for 100 randomly selected clusters whether there are cluster-specific traits (e.g., number of haplotypes, maximum pairwise distance) that can predict whether a barcode cluster will be incongruent with morphology. We identify two such predictors and confirm their effectiveness when applied to the remaining barcode clusters. This leads to the development of explicit rules for picking specimens for validating preliminary species hypotheses based on barcodes. We then compare the barcode clustering results based on different methods, which illustrates once more that integrative methods are needed because COI barcodes alone are insufficiently decisive. Finally, we formalise our system in an algorithm, demonstrating that it is systematic, objective, and effective for our sample.

## Methods

### Sampling

Samples were collected with Townes-style Malaise traps (Townes 1972) at thirty-six sites across Sweden as part of the Swedish Insect Inventory Project (Fig. 1a; Karlsson et al. 2020). A single sample from late spring/early summer 2018 (except for site 46, where the first available sample was from July) was selected from each site (Supplementary Table S1) and specimens from the dipteran family Phoridae were extracted for sequencing. Due to the high number of phorids present (numbering in the hundreds of thousands), only a randomly selected subsample of the specimens was sequenced. The specimens were kept in ethanol at −20-25°C until processing.

**Figure 1.**
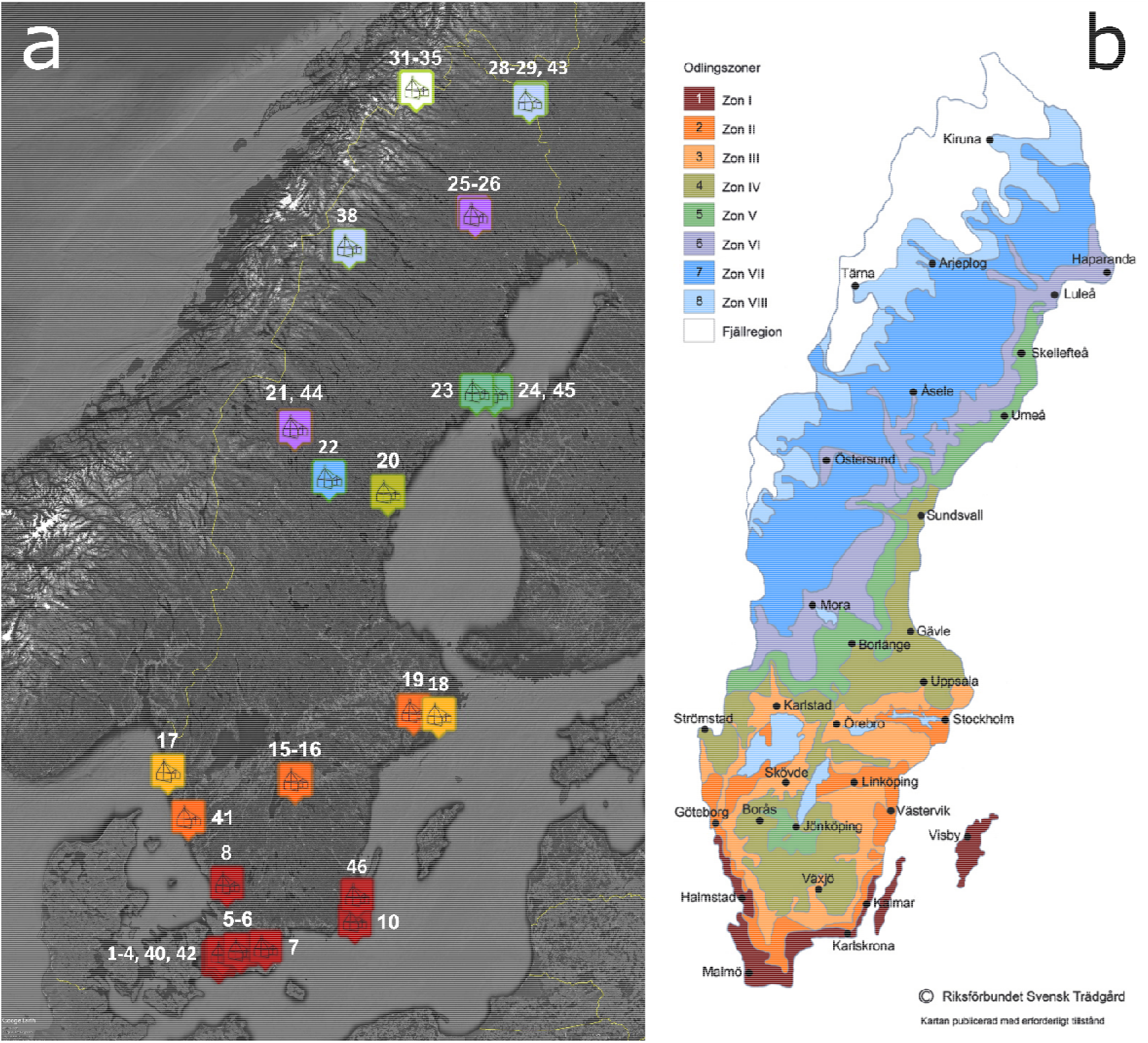
(a) Sites of the Swedish Insect Inventory Project, colour-coded by climatic zones identified by the Swedish Horticultural Society, (b) Climatic zones (odlingszoner) of the Swedish Horticultural Society (Riksförbundet Svensk Trädgård), used with permission.

### DNA Extraction, PCR, and Sequencing

DNA extractions were carried out non-destructively on whole flies using 10 μl of “HotSHOT” solution (Truett et al. 2000, Srivathsan et al. 2019. Incubation was in a thermocycler at 65 °C for 15 min followed by 98 °C for 2 min. A total of 206 96-well plates (19 570 specimens) were extracted. The DNA extracts were used to set up plates of polymerase chain reactions (PCR) to amplify a 313-bp minibarcode fragment of the COI barcoding region using m1COlintF: 5’-GGWACWGGWTGAACWGTWTAYCCYCC-3’ (Leray et al. 2013) and modified jgHCO2198: 50-TANACYTCNGGRTGNCCRAARAAYCA-3 (Geller et al. 2013). Amplifications were conducted with tagged primers and sequenced with Illumina HiSeq 2500 or Oxford Nanopore Technologies (ONT) MinION following the protocols first established in Meier et al. (2016) and then modified for Wang et al. (2018) for Illumina and Srivathsan et al. (2019) for MinION. Note that the most updated lab and bioinformatic methods for MinION barcoding are described in Srivathsan et al. (2021).

PCR reactions contained 4 μl Mastermix from CWBio, 1 μl of 1 mg/ml BSA, 1 μl of 10 μM of each primer, and 1 μl of DNA. PCR conditions were a 5 min initial denaturation at 94 °C followed by 35 cycles of denaturation/annealing/extension (94 °C (1 min)/47 °C (2 min)/72 °C (1 min)), and a final extension at 72 °C (5 min). PCR products were pooled, cleaned, and sequenced in either a lane of HiSeq 2500 (250 bp paired-end sequencing) or a MinION R9.4 flowcell. For MinION, the SQK-LSK109 ligation sequencing kit (Oxford Nanopore Technologies) was used for preparing a library from 200-ng of the pooled and purified PCR products for sequencing. The manufacturer’s instructions were followed except for the use of 1× instead of 0.4× Ampure beads (Beckmann Coulter) because the amplicons in our experiments were short (~391 bp with primers and tags). Illumina libraries were prepared using TruSeq DNA PCR-free kits to obtain 250bp PE sequences using Illumina HiSeq 2500. Illumina sequencing was outsourced. Nanopore sequencing using a MinION sequencer was conducted in house following the description in Srivathsan et al. (2019) and base calling was conducted using Guppy 2.3.5+53a111f (fast basecalling, capacity for high accuracy basecalling was not available at this time).

### Bioinformatics

Demultiplexing and read filtering followed the established protocols in Wang et al. (2018) for Illumina and Srivathsan et al. (2019) for MinION data. Illumina data processing involved merging of paired end reads using PEAR (v 0.9.6) (Zhang et al. 2014). Subsequently, reads were demultiplexed based on unique combinations of tags by an inhouse pipeline that looks for perfect tags while allowing for 2-bp mismatches in primer sequence. For each sample, we retained reads that were longer than 300-bp after primers were removed and the identical reads (reads that only vary in length with terminal bases missing or with extra terminal bases) were merged and counted to identify the dominant sequence, i.e., the sequence with highest count in the dataset. Barcodes were called when (1) the sample contained >=50 reads, and (2) the dominant read had a coverage of over 10×. Lastly, (3) these reads were accepted as the barcode for the specimen if the dominant sequence was at least 5× more common than the sequence with the next-highest abundance. For MinION data, miniBarcoder (Srivathsan et al. 2018, 2019) was used to demultiplex the data. Primers were found using *glsearch36,* and tags extracted allowing for 2 bp errors. Reads with a given tag combination were added into “specimen bins” and those bins having more than 5 reads were processed using multiple sequence alignment via MAFFT. Afterwards, a majority rule consensus barcode was obtained, but only accepted as barcode if it had <1% N (or 4 Ns in case of 313 bp barcodes). A second set of barcodes was obtained from the same bins using the consensus polishing tool RACON, where reads are mapped back onto the original MAFFT consensus barcode using GraphMap and RACON is used to call the consensus sequences (Sović et al. 2016; Vaser et al. 2017). The two sets of barcodes for the same reads were then corrected using amino acid translations that allow for resolving indel errors. Lastly, the AA-corrected sets of barcodes were consolidated as described in Srivathsan et al. (2019). This pipeline has been shown to provide >99.99% accurate DNA barcodes (Srivathsan et al. 2019) and is available at https://github.com/asrivathsan/miniBarcoder.

### Clustering and mOTU estimation

The DNA barcodes were aligned using MAFFT v 7.310 (Katoh and Standley 2013). Many different clustering algorithms for DNA barcodes exist and we here initially used objective clustering (OC, Meier et al. 2006) but later compared the results with other methods. Objective Clustering uses an *a priori* distance threshold to group sequences; cluster members are separated by at least this distance from members of all other clusters, but the maximum distance within a cluster can exceed the clustering threshold (Meier et al. 2006). Initial distance-based mOTU delimitation via objective clustering used a 3% minimum pairwise interspecific threshold done with a new implementation of the clustering algorithm implemented in SpeciesIdentifier that is part of “TaxonDNA” (see Meier et al. 2006: available at github.com/Gaurav/taxondna). Cluster numbers referenced throughout the manuscript refer to these original 3% OC clusters. The clustering results were used to physically sort specimens into putative species in preparation for integrative taxonomy utilising morphology as a second source of data. To visualise the barcode data for each cluster, we prepared median-joining haplotype networks with PopART (Leigh and Bryant 2015). These networks provided a good overview of the number, abundance, and distribution of haplotypes in Sweden. The colour coding reflects the planting regions (“odlingszoner”) recognised by the Swedish Horticultural Society (Fig. 1b, “Zonkarta för odlingzoner”, used with permission) (Riksförbundet Svensk Trädgård 2018). These provide a breakdown of Sweden into eight climatic zones and the alpine zone (“fjällregion”) that coincide with important shifts in the composition of the natural flora of the country. The zones were originally intended to describe the expected success of growing various apple cultivars, but the interpretation has since expanded to cover all cultivated woody plants (trees and bushes grown as ornamental plants or for their fruits or berries) (Riksförbundet Svensk Trädgård 2018).

### Acquisition of morphological data for barcode clusters

Morphological examination of specimens was first conducted in ethanol before some specimens were dissected and slide-mounted in Canada balsam following standard procedures for Phoridae (Disney 2009). Morphological examination relied upon a standard suite of characters and character states described for *Megaselia* (Hartop and Brown 2014; Hartop et al. 2016) and slightly modified for other genera. These traits include a set of characters covering everything from overall gestalt, to setation of the thorax, legs, and frons, to characteristics of the wings and male genitalia. As this method is designed to be efficient, morphological validation did not include more time-consuming morphological methods such as wing measurements or dissection of the genitalia.

### Integration of barcodes and morphology I: Identifying predictors of conflict

One-hundred clusters were randomly selected for in-depth study of congruence between barcode and morphological data. The following rules were applied when selecting specimens for morphological study: (I) Study at least one male for all main haplotypes (= those containing 20% or more of the specimens), (II) study additional specimens within the 3% cluster until no haplotypes remain that are >1 bp away from any checked haplotype, and (III) pick at least one specimen from each horticultural zone represented in the cluster. In most cases, following these procedures was straightforward. For example, Cluster 293 (Fig. 2) has two main haplotypes that each contain >20% of specimens and a singleton that is 2 bp away from the checked haplotypes, meaning at least three specimens must be checked across haplotypes. Additionally, seven zones are represented and a specimen from each zone must be checked, meaning the morphology of at least 7 specimens had to be examined (across both zones and haplotypes). The selection of specimens can be ambiguous because multiple sets of specimens satisfy the stipulated criteria. We then arbitrarily chose one of the potential sets. Additional specimens were studied whenever a barcode cluster was composed of >1 morphospecies. In such cases, we had to determine which haplotypes within the barcode cluster belong to which morphospecies. One barcode cluster contained at least 25 morphospecies (Cluster 101: Fig. 3) and had to be re-clustered at successively lower thresholds (2% and 1%) to help with specimen selection. After reclustering at lower thresholds, the standard checking rules were applied to the sub-clusters.

**Figure 2.**
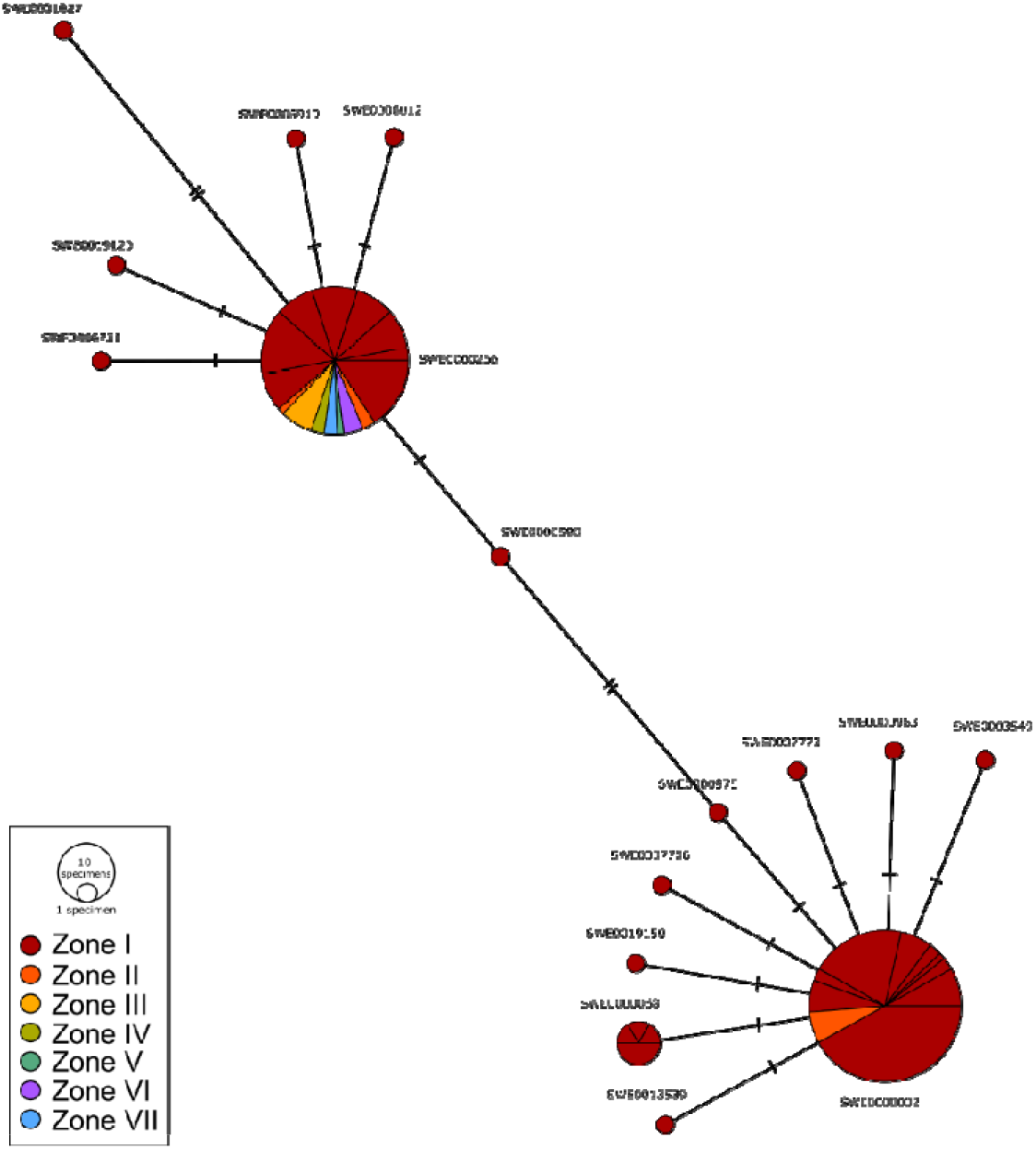
Haplotype network for Cluster 293, colour-coded according to the climatic zones of the Swedish Horticultural Society. Nodes represent each unique haplotype, pie slices of nodes indicate the proportion of specimens from a particular site, node diameters are proportional to the number of specimens the haplotype contains, and the lines connecting the nodes have hash marks corresponding to base pair differences.

**Figure 3.**
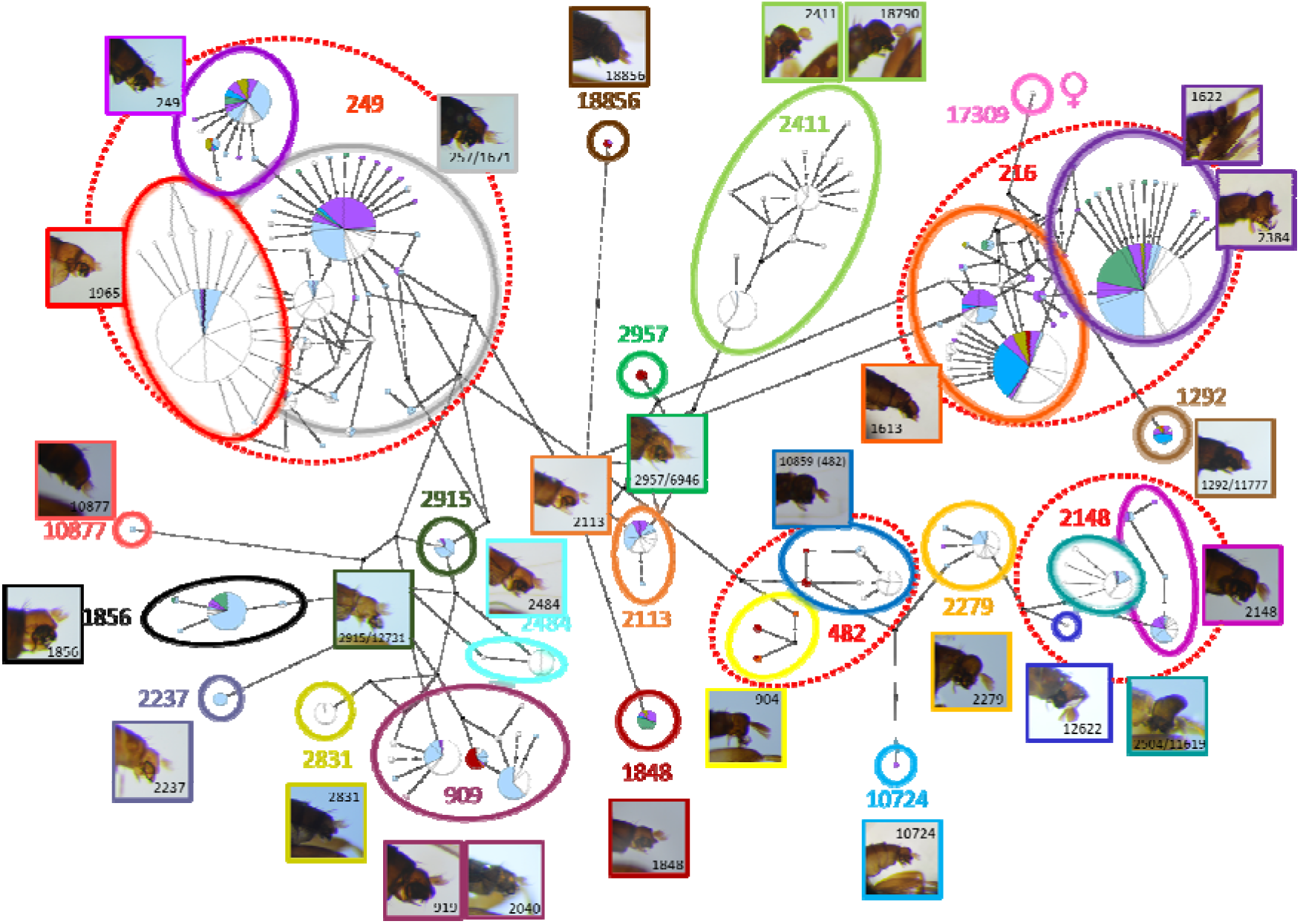
Haplotype network for cluster 101 indicating all morphological species found with male genitalia illustrated (border colours of genitalia figures match morphospecies boundary colours). Morphospecies are equivalent to 1% clusters (indicated by numbers), except in cases where a 1% sub-cluster contained multiple morphospecies, in these cases the 1% cluster is a red dashed line around the morphospecies. For two sub-clusters (216 and 249), the network is too complex to accurately circumscribe morphospecies in this figure. Morphospecies designations for all specimens are in the cluster table available on the project GitHub page.

Note that for all clusters, only males were considered fully informative because much of phorid taxonomy relies on male morphology. Females were treated as congruent if they did not disagree with the males for the cluster regarding key diagnostic characters, but they were not considered sufficiently informative to evaluate distant haplotypes (>1% away) unless they belonged to groups for which female morphology is diagnostic.

To test which cluster-specific properties were most effective at predicting conflict with morphology, we fitted a generalised linear model (glm) with quasibinomial errors to the data obtained for the 100 randomly selected clusters. The response variable was “validated” (whether a cluster was congruent with morphology) and the explanatory variables were six cluster properties: “haplo” (number of haplotypes in a cluster), “spec” (number of specimens in a cluster), “stability” (see below), “max_p” (maximum pairwise distance within a cluster), “zones” (number of geographic zones represented in a cluster), and “sites” (number of sites represented within a cluster).

*Cluster stability* quantified whether a barcode cluster was sensitive to changes in clustering thresholds. This is formalized as class II specimen congruence in Yeo et al. (2020). For a set of clusters at 1%,, that combine to form a single 3% cluster,, the stability value is given by 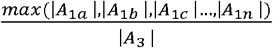, where *A* is set of unique haplotypes and |*A*| is the number of elements in *A*. Simply, this is the ratio between the number of haplotypes contained in the largest 1% cluster that is found within a 3% cluster, divided by the number of haplotypes in that 3% cluster.

A correlation matrix was used to examine collinearity between explanatory variables (corrplot.mixed in R package corrplot) (Supp. Fig. S1) and we then used the Farrar-Glauber test (R package mctest) to detect and remove variables systematically according to the variable inflation factor (VIF), rerunning the model until collinearity was no longer detected and the remaining variable(s) were statistically significant. Due to uncertain species numbers in two clusters designated as “species complexes” (see *Results*), we constructed the model twice: once with these counting as validated clusters corresponding to a single morphospecies, and again with them counting as incongruent (containing multiple species).

### Integration of barcodes and morphology II: Validating predictors of multi-species clusters

After developing predictors of incongruence based on the first 100 clusters, the remaining 215 were used to test the two most important predictors of incongruence; low cluster stability (<1.0) and large intra-cluster p-distances (>/=1.5%). Forty-three of the 215 clusters were identified as PI (Potentially Incongruent) while 172 were non-PI. The 43 largest non-PI were used as control for the 43 PI clusters. These 86 test clusters went through the following validation process: (1) We checked one specimen each for all main (>20% of specimens) haplotypes and (2) a pair of specimens representing the maximum p-distance in the cluster. For the selection of most distant haplotypes, we ignored small differences (1-2 base pairs) in favour of sampling main haplotypes (see *Results*). Using our previous example of Cluster 293 (Fig. 2), the two main haplotypes would be checked, but the haplotypes “haloing” the main haplotypes by 1-2 base pairs would be ignored. If a cluster was found to contain multiple morphospecies during the initial check, additional specimens were studied across the cluster to delimit morphospecies boundaries. For the remaining 129 non-PI clusters, we only checked a pair of specimens representing the maximum p-distance in the cluster.

### Evaluation of the performance of different clustering algorithms and thresholds

We varied the threshold for Objective Clustering and evaluated the performance of three alternative procedures for barcode clustering. The first was Automatic Barcode Gap Discovery (ABGD) (Puillandre et al. 2012), the second Poisson tree process (PTP) (Zhang et al. 2013) and the third Refined Single Linkage (RESL) (Ratnasingham and Hebert 2013). ABGD is distance-based like OC and attempts to find a barcode gap (difference between inter- and intraspecific distances) for each subgroup of sequences based on an iterative application of priors for intraspecific divergence and identification of the first significant gap beyond this divergence. PTP is tree-based and uses branch lengths to estimate transition points between intraspecific and interspecific branching in an input phylogeny, thereby determining which monophyletic clades likely consist of a single species. RESL is used to calculate Barcode Index Numbers (BINs) on the Barcode of Life Data System (BOLD). The underlying algorithm is not public. Instead, new BINs and old BINs are determined or updated monthly by the Canadian Centre for DNA Barcoding.

OC was done at 0.0-5.0% uncorrected *p*-distance thresholds at 0.1% intervals (thresholds between 0.5-4.0% were considered for delimitation comparison, see Results). For ABGD estimation of mOTUs, we used the default range of priors (*P* = 0.0010, *P* = 0.0017, *P* = 0.0028, *P* = 0.0046, *P* = 0.0077, *P* = 0.0129, *P* = 0.0215, *P* = 0.0359, *P* = 0.0599); these priors represent the maximum intraspecific divergence in the first iteration of the algorithm. The slope parameter (-X) was reduced in a stepwise manner (1.5, 1.0, 0.5, 0.1) if the algorithm could not find a partition, as done by Yeo et al. (2020). All other parameters were kept as default. For PTP, we used RAxML v8.4.2 (Stamatakis 2014) to estimate the topology under the GTRGAMMA model. Twenty independent searches were conducted, and the best scoring topology across these searches was retained. mOTUs were then obtained based on the application of the PTP model on this topology, as implemented in the mPTP software (--single --ml mode) (Kapli et al. 2017). We used two indirect methods for evaluating RESL performance for our data, as the algorithm is not public, and the current implementation of the algorithm does not assign new BINs to minibarcodes. We initially identified those haplotypes in our dataset that have a 100% match to haplotypes that are already on BOLD. We then assigned the corresponding BIN numbers to the barcodes for our specimens with these haplotypes. Afterwards, two congruence analyses were carried out. One was based on all our data, but only scoring congruence for specimens with BIN numbers. A second was based on a reanalysis (re-clustering) of only those haplotypes with BIN numbers. Note that both comparisons could be impacted by differences in the underlying data that are used to assign barcodes to BINs.

Congruence between mOTUs and morphospecies was assessed based on match ratio as described in Ahrens et al. (2016). The match ratio is computed as 2 * *N_match_* / (*N_x_* + *N_morph_*), where *N_match_* is the number of completely matching clusters, *N_x_* is the total number of clusters (mOTUs) identified by method *x*, and *N_morph_* is the total number of morphospecies. We also evaluated the performance of clustering methods by determining the number of mOTUs that contained several morphospecies (merged clusters, *N_merged_*), that contained only part of one morphospecies (split clusters, *N_split_*), or the few cases where a method both split and merged members of a single morphospecies *N_split/merged_*. Note that *N_x_* = *N_match_* + *N_merged_* + *N_split+_ N_split/merged_*. Next, we recorded the total number of specimens in each type of cluster. The results were visualised with nVenn (Pérez-Silva et al. 2018) for both optimal (best match to morphospecies) and conservative (above splitting of morphospecies) settings.

To formalise our methods, we created a pairwise distance matrix with MEGA X with default settings (Kumar et al. 2018). This was used as input in the creation of a specimen picking algorithm (Supp. Fig. S2). Using this algorithm, we computed the minimum number of specimens that must be examined morphologically to validate barcode clusters for different methods and settings if the following simplified sampling scheme was applied: (1) determine if the cluster is PI or non-PI (here, based on maximum p-distance only, see discussion), (2) check the main haplotypes and two most distant haplotypes for PI clusters, and two most distant haplotypes only for non-PI clusters. If checked specimens do not belong to the same species, (3) check additional haplotypes until no unchecked haplotypes remain that differ by >1 bp from any checked haplotype. Additionally, all checked haplotypes separated by this distance must belong to the same morphospecies (i.e. there are no unchecked specimens between specimens that belong to two different morphospecies). We also calculated the number of morphospecies (if any) that would be overlooked by this minimal sampling procedure.

### Data and code availability

Data and scripts are available at https://github.com/ronquistlab/taxon-cluster-paper.

## Results

### Barcoding and initial clustering

The 19 570 phorids in the sample yielded 17 902 barcodes in 1 714 haplotypes and 340 clusters at a 3% OC threshold. Of the 340 clusters, 329 were analysed further. Eleven had to be removed because they had been incorrectly pre-sorted to phorids and instead contained different Diptera families (9 clusters totalling 10 specimens), were contamination (a single cluster of 42 specimens that contained a mix of disparate species) or were lost during molecular processing (one singleton cluster). Of the 329 clusters, 14 clusters (406 specimens) could not be rigorously evaluated with morphology because they contained only female specimens, or the most-distant haplotypes were only represented by female specimens. The final analysis thus focused on 17 443 specimens in 315 clusters obtained at a 3% OC threshold.

Of the 315 clusters that were morphologically evaluated, 5.7% (18/315) contained multiple morphospecies. The 18 multi-species clusters contained 18.6% of the species (68/365) and some were among the most abundant in terms of specimen numbers. For example, 19.6% of all specimens were found in the largest incongruent cluster (Cluster 101) while an additional 8.8% of specimens belonged to the second largest cluster (Cluster 68). A total of 7 150 specimens (41% of the total) belonged to clusters that contained more than one morphospecies. All multi-species clusters belonged to the genus *Megaselia*. There were no cases of morphospecies splitting across multiple 3% OC clusters.

### Integration of barcodes and morphology I: Identifying predictors of conflict

Of the 100 clusters randomly selected for the exploratory phase, seven contained multiple morphospecies and two clusters were found to contain morphological variation that was suggestive of species complexes that required more data for resolution. Of the seven incongruent clusters, six split into two species each after morphological examination, while one cluster (Cluster 101: 3 421 specimens) contained at least 25 species (exact species count will require more data to resolve some of the sub-clusters) (Fig. 3). The 7% multispecies clusters (7/100) contained 28% of the species (37/130) and 58.6% of the specimens (3501/5977).

Although the validation procedure involved the examination of many specimens, we noted that the multispecies nature of a cluster was always revealed by the morphology of the two most distant haplotypes. Note that for determining the most distant haplotypes, we exclude “satellite” haplotypes that often “halo” around main haplotypes and differ only by 1-2 bp (see Fig. 2). We examined some of the specimens within these satellite haplotypes, but never found additional morphospecies among them. In the exploratory phase we ensured that specimens from all horticultural zones were examined for all clusters; however, the extra specimens added to satisfy this requirement never resulted in the discovery of additional species.

The exploratory study suggested that the number of specimens (“spec”), the number of haplotypes (“haplo”), the number of collecting sites (“sites”), the number of horticultural zones (“zones”) the maximum p-distance (“max_p”) and the stability to varying clustering thresholds (“stability”) might be correlated with the presence of multiple species within clusters, and all of these were initially included in our model. Three pairs of variables had high collinearity: “max_p” and “stability” (0.77), “haplo” and “spec” (0.99), and “zones” and “sites” (0.79). We systematically removed variables from the model based on the VIF until collinearity was no longer detected, and were left with “spec”, “stability”, and “zones”, but only “stability” was significant (Supplementary Fig. S1). Results of the linear model were the same regardless of whether the two species complexes were classified as validated or not (Supplementary Fig. S1).

Our analysis therefore indicates that only the variable “stability” is a significant in predictor of cluster failure, but the high collinearity (0.77) between “max_p” and “stability” implies that it, too, would be predictive (Supplementary Fig. S1). We subsequently noticed that only 6 of the 7 multispecies clusters were unstable but all of them had high p-distance. This suggested that p-distance should also be used to identify potentially incongruent clusters. Using only the p-distance or the two variables in combination (either/or) ensured that all 7 clusters were identified but it also significantly increased the false positive rate: 22 clusters had high maximum p-distance, but just 12 of these were unstable. Although the combination of the two variables yields a high false-positive rate, we used both criteria for identifying PI clusters among the remaining 215 clusters. More efficient processing could be achieved by dropping one criterion. However, this would result in overlooking the conflict between data sources for some clusters.

### Integration of barcodes and morphology II: Validating remaining clusters and testing predictors for multi-species mOTUs

After in depth study of the first 100 clusters, 215 OC clusters remained (3% threshold). We identified 43 clusters that had signatures of incongruence because of high maximum distances (>1.5%: 14 clusters), instability (stability < 1.0: 2 clusters), or both (27 clusters). Of these PI clusters, a large proportion (26%: 11/43) containing 49% of the species (31 /63) failed morphological validation, and one additional cluster was classified as a “species complex”. All eleven failed clusters had both high maximum *p*-distances and instability, while the species complex was identified by maximum p-distance only. To determine whether 26% failure is an unexpectedly large proportion of failing clusters, we used the largest 43 non-PI clusters as control. The largest non-PI clusters were used to best match cluster sizes for PI (mean=152 specimens) and non-PI clusters (mean=95). All 43 non-PI clusters passed morphological validation as did the remaining 129 smaller, non-PI clusters.

The total number of specimens examined for this study was 1 039, or 5.8% of the total. This includes the specimens for the more extensively sampled first 100 clusters. It also includes some female specimens that represented haplotypes for which a male was also examined. Without these additional specimens, the number of specimens needed for validation of 315 3% OC clusters would have been 915 (5.1% of total). The optimised number of specimens that needs to be studied is 861 (4.8% of the total) when using OC 1.3-1.5% or ABGD 0.0077 as clustering thresholds. This is an average of 2.3 specimens per species.

### Impact of barcode clustering algorithms on results

Our morphological study suggested the presence of 365 morphospecies and a match ratio of 0.871 for 3% OC clusters (Fig. 4, Table 1). All incongruence between 3% OC clusters and morphology was due to lumping. Lowering the threshold to 1.6-1.7% maximised the match ratio for OC at 0.897 (Fig. 4, Table 1). ABGD’s highest congruence was the same (0.897) and observed when the prior of intraspecific divergence (p) was set to 0.0077 (Fig. 4, Table 1). PTP fared less well, with a match ratio of 0.841 (Fig. 4, Table 1). Regarding RESL, 277 of the 1714 haplotypes in our dataset had 100% matches to publicly available sequences with BIN designations. These 277 haplotypes represented 50% (186) of our morphospecies in 172 BINs. Of the 186 morphospecies matched to BINs, 162 (86%) were congruent with BINs. These 277 haplotypes were also subjected to re-clustering with several algorithms. We compared the results for both optimal (highest match ratio) and conservative (above cluster splitting) threshold settings for OC and ABGD. BIN designations again matched 86% of morphospecies. This was better than OC at the conservative threshold of 3% (76% of morphospecies correctly delimited), but worse than OC at the optimal 1.7% setting (89% of morphospecies correctly delimited). PTP performed poorer for this reduced subset (74%), and ABGD had the lowest level of congruence (correctly delimiting 72% of morphospecies at p=0.0077 but only 51% at p=0.0215).

**Figure 4.**
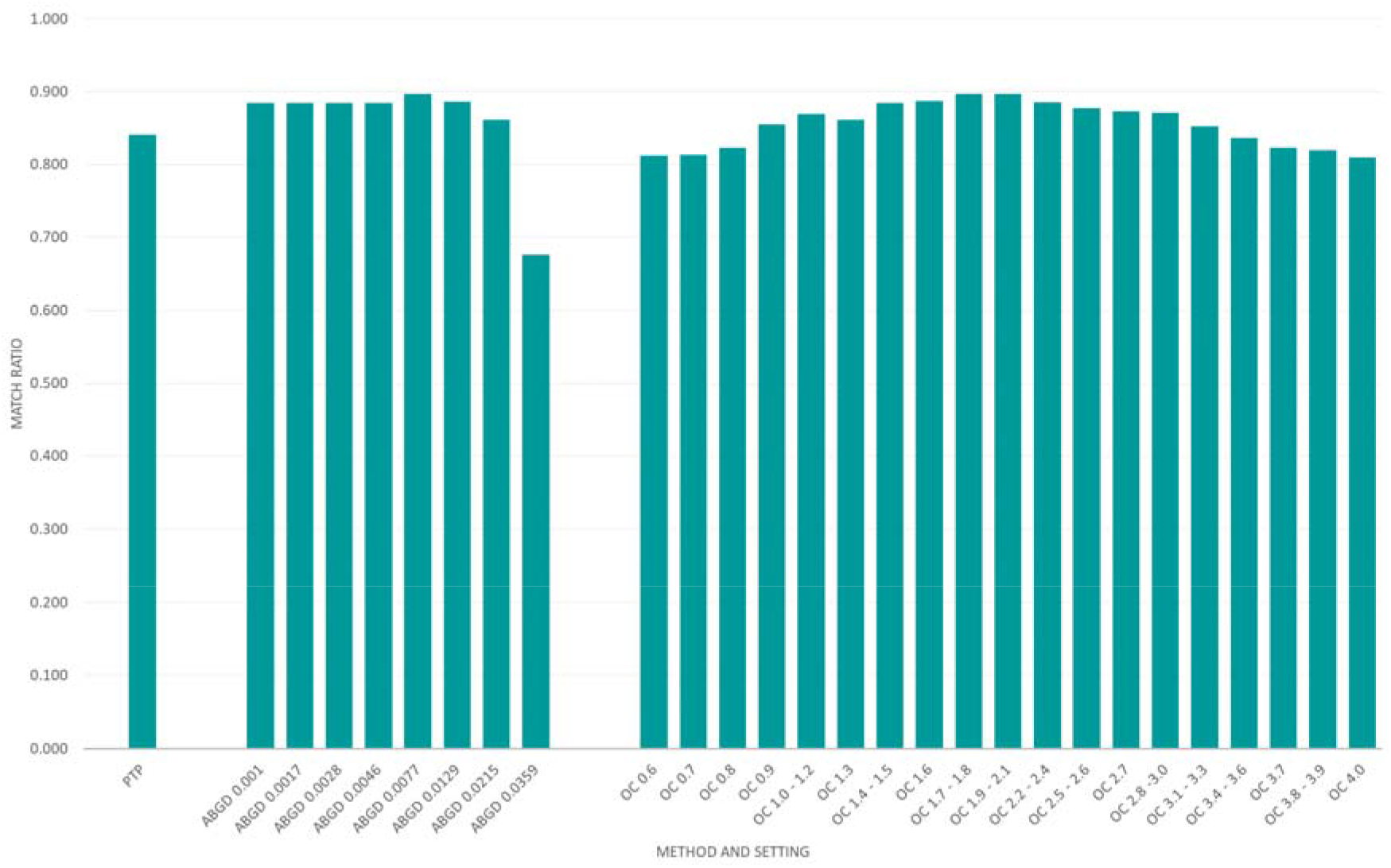
Match ratios for PTP, ABGD (all priors) and OC (all thresholds) versus morphology across methods and settings.

**Table 1.** Summary of mOTU delimitations across methods and settings. Optimal match ratios are highlighted in green.

|  | # of mOTUs | # mOTUs congruent with morphology | # of morphospecies lumped | # of morphospecies split | # of morphospecies split + lumped | # units molecular delimitation only | # units morphospecies only | Match Ratio | Specimens to check |
| --- | --- | --- | --- | --- | --- | --- | --- | --- | --- |
| PTP | 394 | 320 | 21 | 25 | 1 | 74 | 47 | 0.841 | 880 |
| ABGD 0.001 | 370 | 326 | 26 | 15 | 0 | 44 | 41 | 0.885 | 861 |
| ABGD 0.0017 | 370 | 326 | 26 | 15 | 0 | 44 | 41 | 0.885 | 861 |
| ABGD 0.0028 | 370 | 326 | 26 | 15 | 0 | 44 | 41 | 0.885 | 861 |
| ABGD 0.0046 | 370 | 326 | 26 | 15 | 0 | 44 | 41 | 0.885 | 861 |
| ABGD 0.0077 | 362 | 327 | 29 | 11 | 0 | 35 | 40 | 0.897 | 876 |
| ABGD 0.0129 | 337 | 312 | 50 | 5 | 0 | 25 | 55 | 0.886 | 898 |
| ABGD 0.0215 | 311 | 292 | 75 | 0 | 0 | 19 | 75 | 0.861 | 915 |
| ABGD 0.0359 | 207 | 194 | 173 | 0 | 0 | 13 | 173 | 0.676 | 1061 |
| OC 0.0 - 0.2 | 1714 | 126 | 0 | 241 | 0 | 1588 | 241 | 0.121 | 1714 |
| OC 0.3 | 552 | 257 | 0 | 110 | 0 | 295 | 110 | 0.559 | 902 |
| OC 0.4 - 0.5 | 542 | 261 | 0 | 106 | 0 | 281 | 106 | 0.574 | 895 |
| OC 0.6 | 424 | 321 | 4 | 41 | 1 | 103 | 46 | 0.812 | 863 |
| OC 0.7 | 423 | 321 | 4 | 41 | 1 | 102 | 46 | 0.813 | 863 |
| OC 0.8 | 418 | 323 | 4 | 39 | 1 | 95 | 44 | 0.823 | 860 |
| OC 0.9 | 396 | 326 | 12 | 29 | 0 | 70 | 41 | 0.855 | 865 |
| OC 1.0 - 1.2 | 390 | 329 | 13 | 25 | 0 | 61 | 38 | 0.869 | 863 |
| OC 1.3 | 390 | 326 | 26 | 15 | 0 | 64 | 41 | 0.861 | 861 |
| OC 1.4 - 1.5 | 370 | 326 | 26 | 15 | 0 | 44 | 41 | 0.885 | 861 |
| OC 1.6 | 370 | 327 | 29 | 11 | 0 | 43 | 40 | 0.887 | 876 |
| OC 1.7 - 1.8 | 362 | 327 | 29 | 11 | 0 | 35 | 40 | 0.897 | 876 |
| OC 1.9 - 2.1 | 347 | 320 | 41 | 6 | 0 | 27 | 47 | 0.896 | 898 |
| OC 2.2 - 2.4 | 336 | 311 | 51 | 5 | 0 | 25 | 56 | 0.885 | 898 |
| OC 2.5 - 2.6 | 324 | 303 | 61 | 3 | 0 | 21 | 64 | 0.877 | 916 |
| OC 2.7 | 323 | 301 | 63 | 3 | 0 | 22 | 66 | 0.872 | 916 |
| OC 2.8 - 3.0 | 315 | 297 | 70 | 0 | 0 | 18 | 70 | 0.871 | 915 |
| OC 3.1 - 3.3 | 311 | 289 | 78 | 0 | 0 | 22 | 78 | 0.853 | 914 |
| OC 3.4 - 3.6 | 305 | 281 | 86 | 0 | 0 | 24 | 86 | 0.836 | 912 |
| OC 3.7 | 299 | 274 | 93 | 0 | 0 | 25 | 93 | 0.823 | 912 |
| OC 3.8 - 3.9 | 297 | 272 | 95 | 0 | 0 | 25 | 95 | 0.819 | 912 |
| OC 4.0 | 293 | 267 | 100 | 0 | 0 | 26 | 100 | 0.809 | 911 |
| OC 4.1 - 4.2 | 292 | 266 | 101 | 0 | 0 | 26 | 101 | 0.807 | 911 |
| OC 4.3 | 284 | 257 | 110 | 0 | 0 | 27 | 110 | 0.790 | 919 |
| OC 4.4 | 283 | 257 | 110 | 0 | 0 | 26 | 110 | 0.791 | 956 |
| OC 4.5 | 281 | 253 | 114 | 0 | 0 | 28 | 114 | 0.781 | 957 |
| OC 4.6 | 272 | 241 | 126 | 0 | 0 | 31 | 126 | 0.754 | 951 |
| OC 4.7 - 4.8 | 269 | 237 | 130 | 0 | 0 | 32 | 130 | 0.745 | 954 |
| OC 4.9 | 256 | 226 | 141 | 0 | 0 | 30 | 141 | 0.726 | 970 |
| OC 5.0 | 254 | 225 | 142 | 0 | 0 | 29 | 142 | 0.725 | 975 |

We then examined whether morphospecies were split, lumped, or split+lumped across different methods and settings (Fig. 5, Table. 1). This revealed that OC starts lumping morphospecies at 0.6% and stops splitting morphospecies at 2.8%, while ABGD lumps morphospecies across all priors, and stops splitting at p=0.0215 (Fig. 5, Table. 1). PTP both splits and lumps morphospecies (Table 1). There were very few cases where a method both split and lumped a single morphospecies, but this was the case for one morphospecies using PTP and at OC thresholds 0.7-0.9% (Fig. 5, Table. 1). RESL split two of the clusters designated as “species complexes” in our analysis but lumped 22 morphospecies into BINs (from OC clusters 68, 79, 91, 101, 103). The BIN composition of the complex Cluster 101 could be assessed for 16 morphospecies (Fig. 6). Based on this partial representation, RESL lumps eight of the morphospecies from Cluster 101 into a single BIN (BOLD-AAG3235, shown in red) and another three into a second BIN (BOLD-AAL9067).

**Figure 5.**
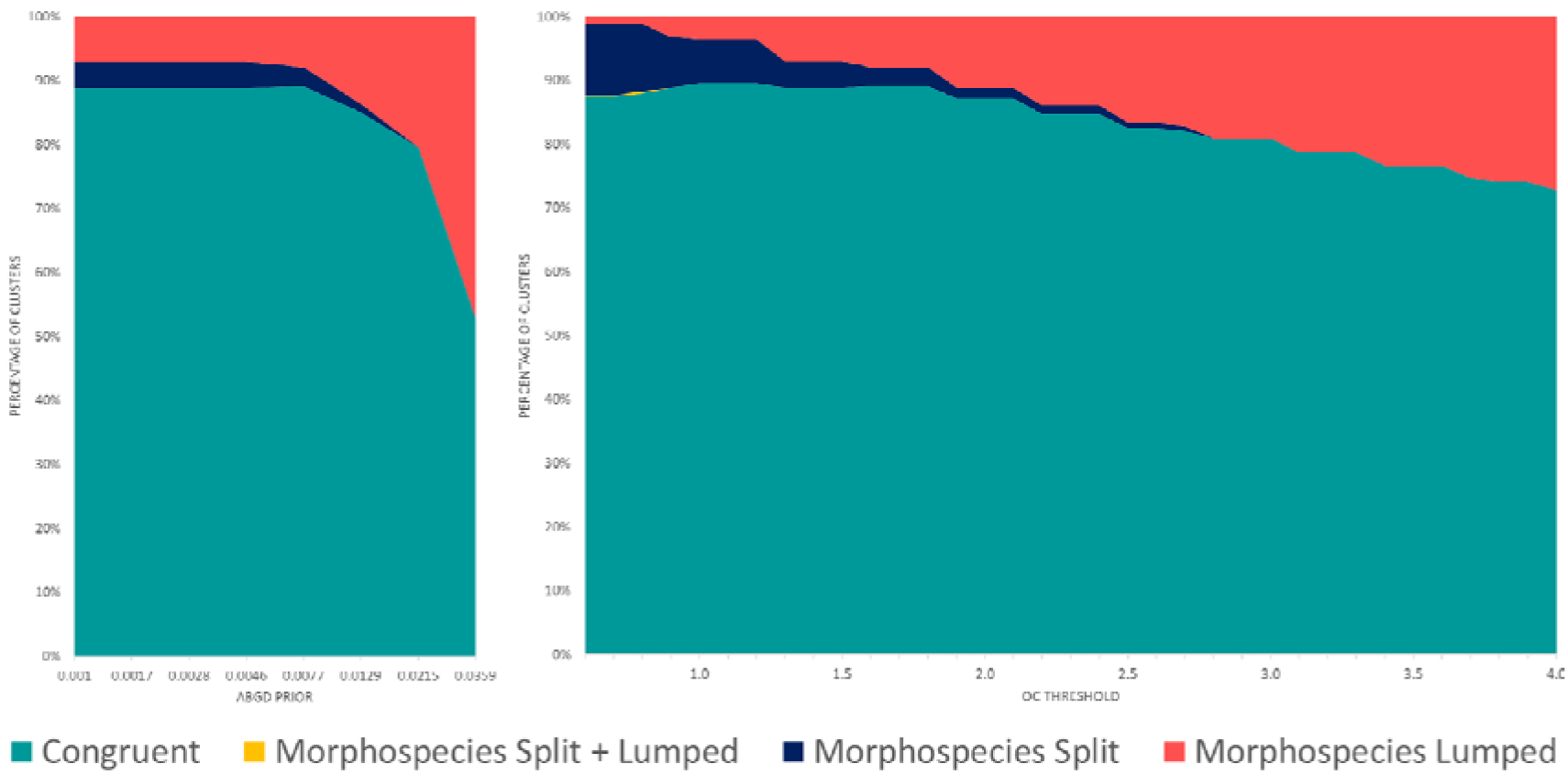
The splitting and lumping of morphological clusters with ABGD (left) and OC (right) across settings.

**Figure 6.**
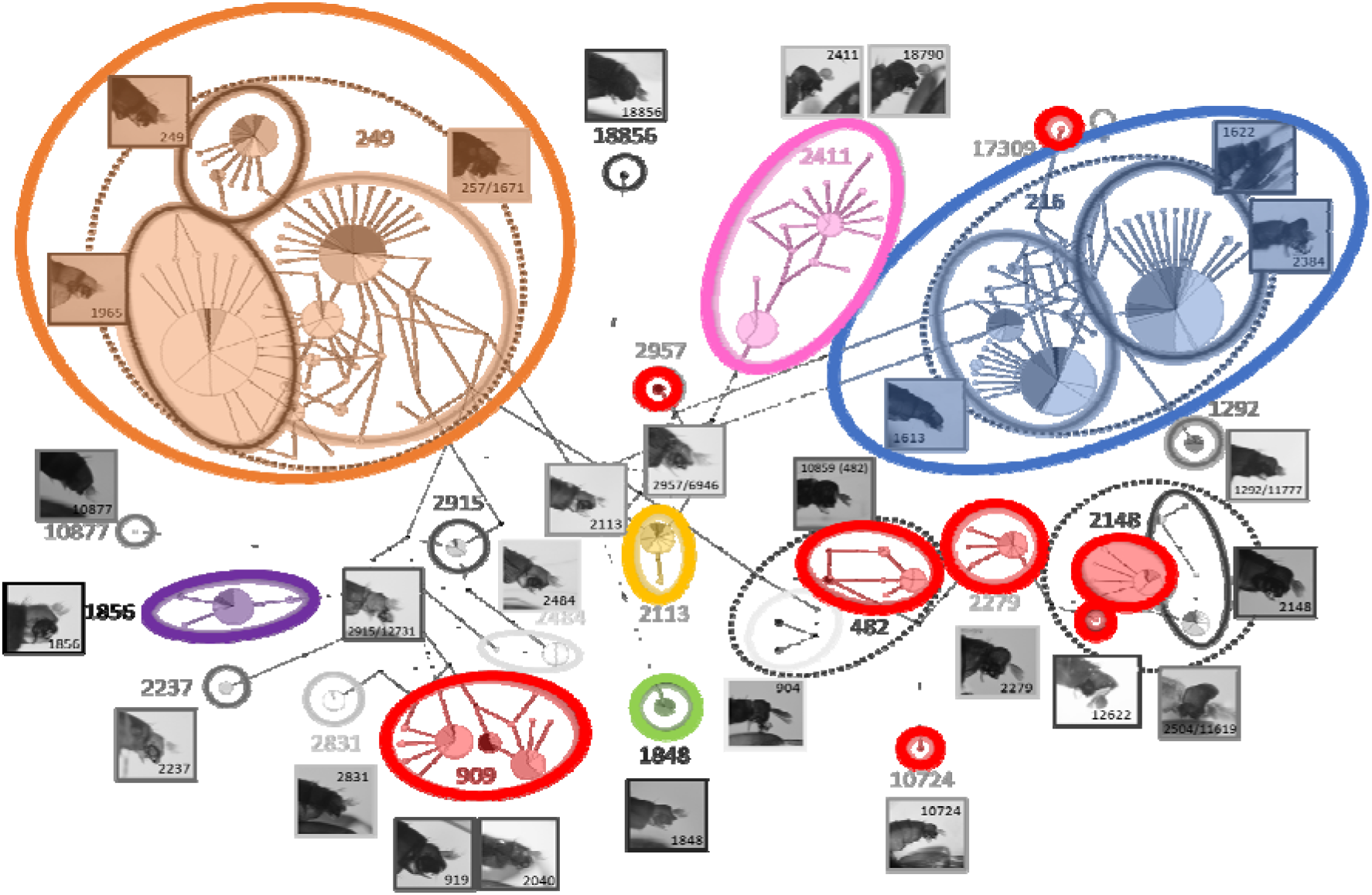
BIN designations of the 16 morphospecies of Cluster 101 for which we found a 100% match (to at least one specimen) in BOLD.

Figure 7 illustrates congruence between methods for optimal (OC 1.7, ABGD p=0.0077) and conservative settings (OC 3.0%, ABGD p=0.0215). At optimal settings, 313 clusters were the same across methods and ABGD and OC results were 100% congruent. PTP was an outlier, as it tended to split species compared to the other methods. Morphology had several dozen outlier species that were different from what was inferred with all molecular methods. This was due to multiple species clustered by ABGD and OC at conservative settings (including the 25 species in Cluster 101), while at optimal settings the difference was due to fewer lumped species, but additional split species. We clustered across a wide range (0-5.0%) of OC thresholds to assess the structure of variation in our dataset. We would not consider thresholds lower that 0.5 or above 4.0% to be appropriate for initial delimitations in most groups unless there is evidence for large genetic variation within species. Narrowing the range of thresholds accordingly, the number of clusters across methods and settings varies from 424 (0.6% OC) to 207 (p=0.0359 ABGD) (Fig. 8).

**Figure 7.**
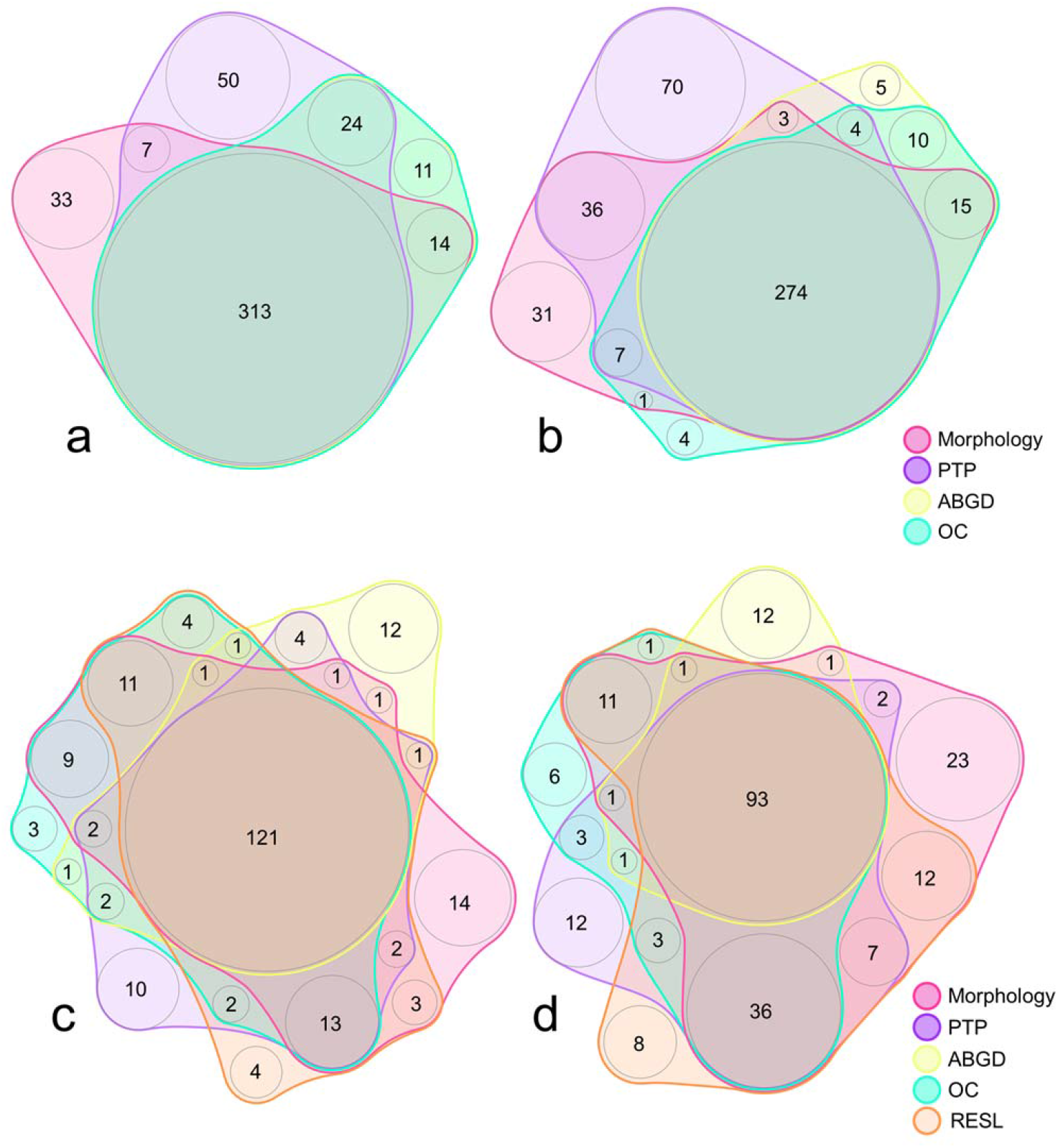
An illustration of the congruence between morphology, PTP, ABGD, and OC methods with (a) optimal settings (ABGD P=0.0077, OC 1.7%) and (b) conservative settings (ABGD P=0.0215, OC 3.0%) and between morphology, PTP, ABGD, OC and RESL methods with (c) optimal settings (ABGD P=0.0077, OC 1.7%) and (d) conservative settings (ABGD P=0.0215, OC 3.0%).

**Figure 8.**
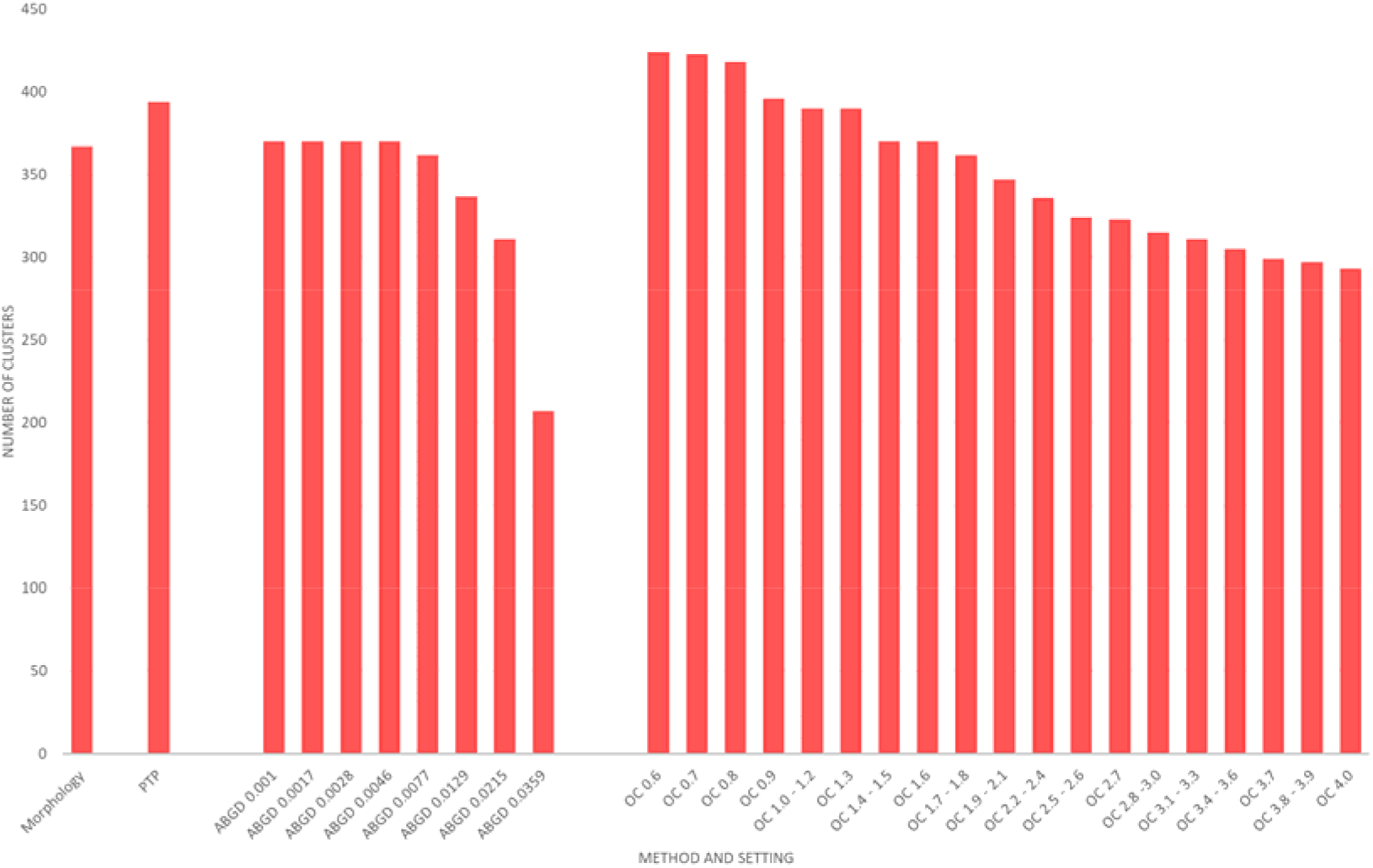
The number of morphospecies, and clusters across settings with PTP, ABGD, and OC. OC is plotted without 0-0.5% thresholds where 1-2 bp differences between haplotypes greatly inflated cluster numbers.

## Discussion

The goal of Large-scale Integrative Taxonomy (LIT) is to transform the study of dark taxa that are currently neglected because traditional methods are unsuitable for samples containing thousands of specimens and hundreds of species. LIT has the potential to enable biodiversity discovery initiatives to take on the challenge that these taxa pose. In the future, new samples of dark taxa could be imaged and/or NGS-barcoded upon arrival. Afterwards, they can be physically sorted to putative species based on barcoding or imaging data. Most of this initial work can be rapid and inexpensive because barcoding and imaging are suitable for automation. Scientists at these institutions will then work only on pre-sorted material, focusing on clusters and specimens that have a high probability of being incongruent with other data sources. These clusters and specimens are identified by a selection algorithm that uses high intra-cluster variation and shallow (below clustering threshold) intra-cluster splits to identify specimens critical for validating barcode clusters or resolving conflict. Morphological or nuclear data can then be obtained for only those selected specimens. Opting to use nuclear data for validation would allow institutions to cover orphaned dark taxa for which the world lacks taxonomic expertise. Using LIT, clusters would be covered by at least two types of data that can be summarised automatically in preparation for species (re)description. Note that LIT is likely to be less costly than many recently funded collection digitisation initiatives. Indeed, LIT is a particularly efficient digitisation initiative, as the imaging of new samples is easier because they are in a more uniform and accessible state than most collection specimens. Furthermore, all locality information for such samples tends to be digitised.

Any large-scale integrative taxonomy initiative must overcome the conundrum of how to accelerate biodiversity discovery based on at least two data sources when most traditional studies only use one and are still too slow. We agree with Puillandre et. al. (2012) that this can be achieved by starting with one type of data that can be acquired rapidly and inexpensively. The second type of data can then be more expensive or time consuming as it will only be obtained for a small subset of specimens. The cumulative effort of acquiring both sets of data can then take a fraction of the time that would be needed using traditional methods. However, for LIT to work, there must be a quantitative and systematic way to flag the clusters and specimens that are most likely to be incongruent with morphology.

We tested several barcode cluster properties and found that cluster stability is the most important predictor for incongruence, but we also used maximum *p*-distance because some incongruent clusters were only identified by this variable. Cluster instability is indicative of areas where multiple species are separated by shallow (below clustering thresholds) splits, while maximum p-distance identifies clusters with unusually high variation. Observing such shallow splits is not surprising because few evolutionary biologists doubt that there are cases of recent and rapid divergences, and these are the areas where barcodes are known to be most likely to fail (Puillandre et al. 2012; Ratnasingham and Hebert 2013; Zhang et al. 2013). We here quantify cluster stability by testing whether cluster membership changes when the clustering threshold is modified. This is suitable for objective clustering but an alternative way to identify unstable clusters would be to inspect the longest branch length on a median-joining network for each cluster (as in the haplotype networks) (see Supp. Fig. S2 for how this might be incorporated to the specimen checking algorithm). High within-cluster distances may be indicative of two species or old lineages within one species that have acquired large amounts of genetic variation but have failed to speciate. Overall, it is not surprising that these cluster properties are associated with conflict, but it was surprising to find that they are so effective at predicting it. Over one-quarter of clusters flagged by the predictors failed validation with morphology, while none of the control, non-PI clusters contained more than one morphospecies. Of course, this result will have to be tested for more taxa and larger samples sizes before a more general use of these predictors can be advocated.

Identifying clusters that are likely in need of refinement is only the first step. LIT also needs rules for selecting those specimens that should be studied using a second type of data (e.g., nuclear genes, morphology). In this context, it is important to minimise the number of specimens examined while ensuring that clusters containing multiple species are reliably discovered. Our final LIT protocol includes specimen-picking recommendations (Fig. 9) that are formalised as an algorithm that is available from the project’s Github repository (Supp. Fig. S2). The basic recommendation is simple and common sense, but our large-scale study shows that it is also effective. All non-singleton clusters are designated as either PI or non-PI (although singleton clusters could be designated PI based on stability if examined at an increased, rather than decreased, threshold). PI clusters are then validated by sampling the main and most distant haplotypes, while the verification of non-PI clusters is based on a pair of specimens representing the most distant haplotypes. Based on our data clustered at 3%, this protocol will reliably distinguish clusters that are congruent from those that are incongruent. Resolving the incongruent clusters then requires more in-depth examination. For our data, we demonstrate that only ca. 5% of all specimens (i.e., 2.5 specimens per species; 915 specimens in total) need to be checked to integrate barcode data with morphology. This number is required when clustering barcodes at 3% thresholds, but it is quite close to the theoretical minimum of 640 specimens (1.7 per species) that could achieved by checking just two specimens for each cluster regardless of PI designation, excluding singletons for which the only specimen would be checked.

**Figure 9.**
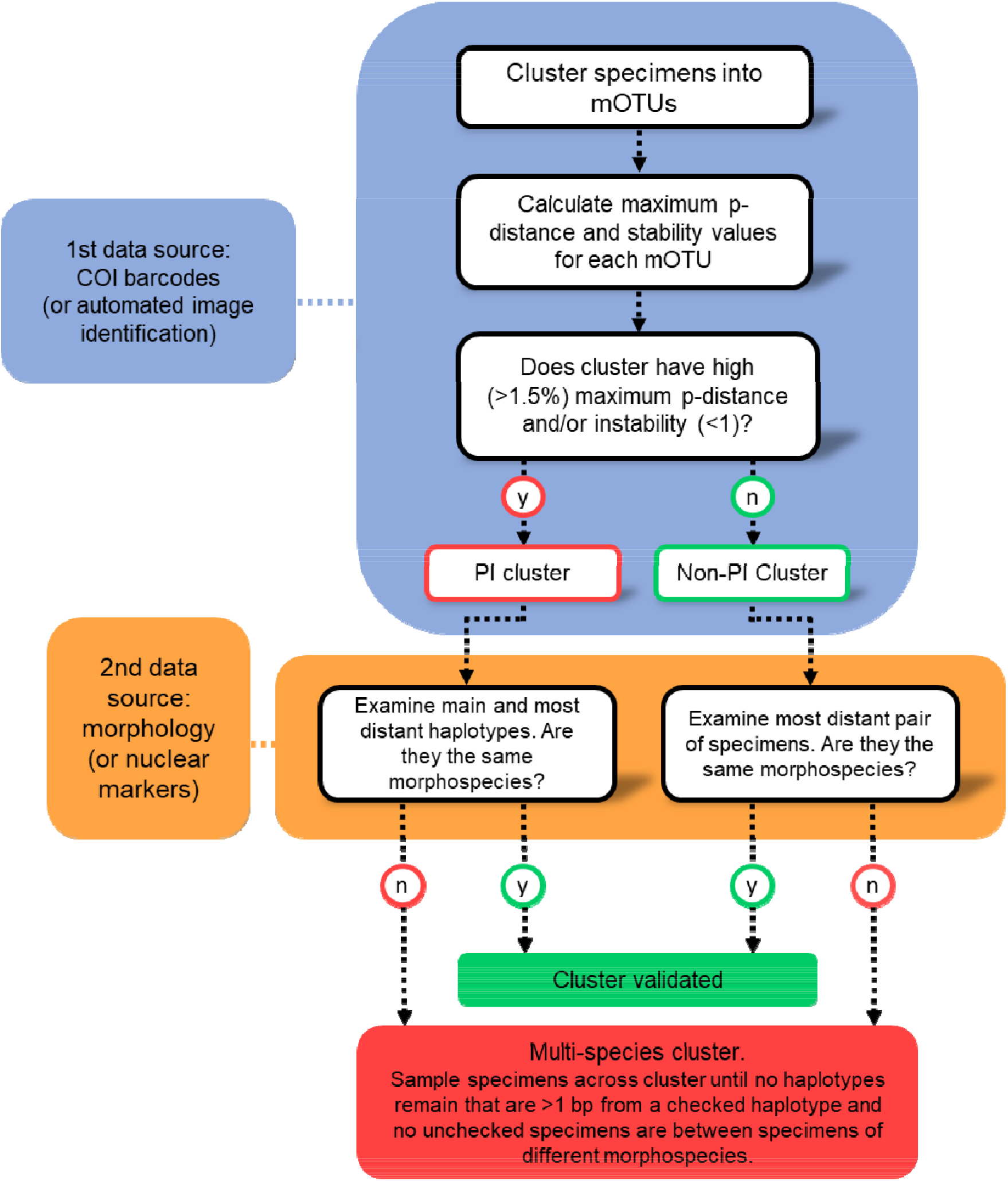
Final LIT protocol. LIT has also been formalised with an algorithm (see Supplementary Figure S2).

The LIT protocol proposed here includes checking main haplotypes for PI clusters because multiple morphospecies can be intermixed within a closely related network (e.g., “subcluster 216” in Cluster 101, Fig. 3). Checking specimens from main haplotypes ensures that the cluster variation observed in a large proportion of specimens is covered. For example, Cluster 293 (Fig. 2) has 163 specimens. Checking only the extreme haplotypes would reveal cluster failure, but the two checked specimens would represent the haplotypes of just 1.2% of the specimens (two singleton haplotypes). Including specimens from each of the two main haplotypes raises the percentage of specimens represented by the haplotype variation that has been checked to 88.3%. Therefore, LIT protocol recommends using the main + distant haplotype checking procedures for PI clusters, and it may be prudent to also check all main haplotypes for large non-PI clusters. Specimen selection can be aided by haplotype networks. These networks allow for a quick visual overview of the structure and patterns of variation within a cluster. Specialists working with the LIT system will find that the use of haplotype networks will facilitate a comprehensive understanding of the molecular variation in their taxa.

One major task when conducting a revision based on 18 000 specimens is specimen handling. We recommend physically sorting all sequenced specimens into vials by haplotype + sample, and then grouping vials together by cluster. Such physical sorting will facilitate morphological work as all specimens will be easily accessible. Fortunately, it should be possible to develop automated ways to physically pre-sort specimens based on barcodes given that the vouchers will be in grid-like microplates. Similarly, distinguishing sexes via AI-assisted image processing should be feasible. We here used scuttle flies from a temperate zone where communities are dominated by species that require males for morphological identification. In the tropics, however, there is a greater abundance of species that are more readily identifiable through examination of female specimens (Borgmeier 1962). In these species, highly distinctive and species-specific female abdominal structures suggest that sexual role reversal likely takes place (Brown and Porras 2015) and therefore it will not always be appropriate for studies examining phorids to eliminate female-only clusters from morphological validation as we did here. We chose to barcode both males and females to make the greatest amount of information available for future uses (associating males and females, increasing abundance and distribution data, etc.).

Our study used data from ~18 000 specimens of scuttle flies (Diptera: Phoridae). Phoridae is a classic example of a dark taxon; i.e., a specimen- and species-rich group where species discovery and identification are wanting. In terms of abundance, the family comprises ca. 10% of the total catch in Malaise trap samples from Sweden (Karlsson et al. 2020). In terms of diversity, it is the genus *Megaselia* that renders Phoridae a dark taxon – none of the other ca. 270 genera come close to the richness of this genus, and most are comparatively well explored. At present, *Megaselia* contains approximately 1 700 described species, but the worldwide fauna may be two orders of magnitude larger (Srivathsan et al. 2019). LIT will have to be tested and adapted to other taxa to optimise the number of specimens to examine. For example, we here used a 3% threshold for initial clustering, but this threshold may not be optimal for all taxa, datasets, or workflows. High thresholds result in fewer clusters to check, but more clusters will fail the congruence test and require checking of main haplotypes in addition to a distant pair. Additionally, high clustering thresholds will result in higher numbers of lumped species, requiring more morphological work to identify the boundaries of morphospecies within the clusters. On the other hand, low thresholds result in morphospecies splitting (Fig. 5, Table 1). Such splitting necessitates careful checking both between and within clusters, which is laborious and time-consuming. An optimal strategy may be to set the threshold just high enough to avoid any splitting of morphospecies, OC at 3% was such a setting. PTP is probably not a good first clustering strategy because it both splits and lumps species.

Other parameters used in LIT also need adjustment to specific taxa and datasets. For example, we here used a 1.5% maximum p-distance or 1 to 3% stability threshold to identify potentially incongruent (PI) clusters, but the corresponding values may be different for other studies. Fortunately, our study reveals that such optimisation can be made based on a small number of randomly chosen clusters and does not increase the workload very significantly. We here had to study only 200 additional specimens to optimize the PI parameters and we predict that fine tuning of LIT for other taxa will require even fewer additional specimens. One way to lower the workload is to redefine “main haplotype” more or less conservatively based on specialist preference. Setting the threshold for main to 10%, one may have check up to 10 specimens per cluster (if a cluster were perfectly spread across 10 haplotypes), while a 20% cut-off would require checking only 5. If too many clusters are revealed to be potentially incongruent, one can increase the efficiency of LIT by using more “lenient thresholds” which may then result in a moderate number of multi-species clusters being overlooked. For example, for our phorid data using the “large p-distance criterion” designated 24 clusters as PI, but in the end this criterion only found one additional incongruent cluster and one species complex.

### Evaluation of a range of mOTU clustering methods

There are many algorithms for clustering barcodes. Most methods include the disclaimer that they should not be used to delimit species without consulting other evidence (Puillandre et al. 2012; Ratnasingham and Hebert 2013; Zhang et al. 2013), but it is not uncommon to see molecular clades equated to species without, or with unexplained, processes of validation. Our study confirms that none of the sequence clustering methods tested (OC, ABGD, PTP, RESL) accurately delimit morphospecies across taxa with disparate evolutionary processes. Adaptive algorithms, such as ABGD and RESL, should be able to accommodate biological variation across subgroups better than methods based on fixed thresholds, but surprisingly our results indicate that a simple method using fixed thresholds (OC) does as well or better than adaptive methods. The tree-based method (PTP) did worst of the tested methods, possibly because the analysis was based on minibarcodes, with limited information about phylogenetic relationships and the boundaries between within-species and between-species tree structure. Tree-based methods should be tested with longer sequences and multiple markers. Such data are also required for other methods that are rooted in the statistical analysis of multispecies coalescent and similar models. LIT will also be relevant for these methods because it predicts for which specimens/haplotypes multiple markers should be collected. After all, for hyperdiverse taxa such data collection will always have to be restricted to subsamples.

Comparisons between methods revealed that there was often better congruence between molecular clusters obtained with different methods than between those clusters and the results obtained with morphology. For example, at optimal thresholds, OC and ABGD produced the same clusters across the dataset, but morphology has an additional 40 morphospecies (Fig. 7a). Similarly, in the data subset used to evaluate RESL, morphology has 14 morphospecies that were not delimited by any of the molecular methods (Fig. 7c). Should we take these examples as evidence that the morphological evidence is misleading? This would be perilous given what is known about barcodes. Even the best algorithms will not be able to accurately delimit species if speciation has left little to no trace in the COI data, as is expected for recently evolved species. For example, PTP was introduced with the “fundamental assumption…that the number of substitutions between species is significantly higher than the number of substitutions within species” (Zhang et al. 2013). Similarly, with ABGD, “the lower the speciation rate, the better the performance of the method” (Puillandre et al. 2012) and RESL carries the warning “closely related species…will be overlooked because of their low sequence divergence” (Ratnasingham and Hebert 2013). Dense specimen sampling may reveal near-continuous sequence variance across thousands of specimens (see Cluster 101) so that any algorithm will struggle to find accurate solutions. Our dataset suggests that such clades of closely related species may be concentrated in some groups. Fortunately, in this study, many species-level taxa that could not be delimited by any of the algorithms applied to barcodes were distinct based on morphological differences (e.g., Cluster 101, Fig. 3).

### The Future of LIT

To evaluate the future utility of LIT, we must first test this system on many other taxa and a wide variety of sample densities. We must determine how LIT protocol can be modified with increased sampling, as the effectiveness of DNA barcoding for delimiting species is dependent on both the depth of sampling and the breadth of geographic scale (Bergsten et al. 2012; Huemer et al. 2014; Ahrens et al. 2016). Datasets that are shallowly sampled or of limited geographic scope can often appear decisive even in species-rich environments like the tropics (Hajibabaei et al. 2006; Smith et al. 2008), but with ever-expanding datasets the complex process of species delimitation (Sites and Marshall 2004; De Queiroz 2007; Wiens 2007) will only be further complicated. As the accumulation of sequence data accelerates, the shortcomings of DNA barcodes (Will et al. 2005; Meier et al. 2006, 2008; Cesari et al. 2011) will only become more evident, and the need for the integration of several types of data will become more apparent (Dayrat 2005; Puillandre et al. 2012).

A true test of LIT thus requires more sampling. Most species in our study were separated by 3% or more but as sampling increases, genetic distances between species will decrease and eventually barcodes may not separate closely related species. In our study using 313 bp minibarcodes, species separated by 3% have a 98.5% chance of being separated by at least 3 bp. However, if species are separated by just 1% genetic distance on average there is a 4.3% chance they would have identical 313 bp barcodes and an 18% chance of the barcodes being separated by just a single bp, ignoring any variation within species. This makes it clear that reducing distances between species may become problematic rather quickly. Even within our dataset, we had some species (such as in Cluster 101, Fig. 3) separated by just 0.6% (2 bp). Such small genetic differences between morphospecies suggest relying on delimitation with barcodes alone will lead to widespread taxonomic error, and such errors will only increase over time (Meier et al. 2021). This is not a problem unique to barcodes, it is a problem with relying on any single data source – morphology is also likely to fail in cases of closely related species. Until we have better datasets with worldwide sampling for many groups, we will not be able to confidently address these issues with LIT. We expect that the situation will be messy, but we also expect many clusters to remain stable.

One strategy for managing difficult datasets is lowering clustering thresholds incrementally as a dataset grows and the distances between clusters shrink. Testing this empirically will require much larger datasets than we have now. If future studies indicate an abundance of fusions, a potential remedy might be to sample more specimens for the second, or add a third data source. COI barcodes could be supplemented not only with morphology, but also with nuclear markers, life history, or ecological data (Puillandre et al., 2012). The standardisation of nuclear markers (Eberle et al. 2020) will be essential to integrating these data and is likely a realistic, long-term approach. Integration of multiple data sources could be multi-layered, e.g., initial clustering with COI, followed by a preliminary validation with morphology, with independent markers added only to those clusters that need further resolution. These data would then also allow for the application of coalescence methods for species delimitation based on the specimens singled out for deeper sequencing.

In this study, we used mini-barcodes (313 bp), as they are easily obtained on short-read platforms such as Illumina and have been shown to perform comparably to full-length barcodes for the identification and delimitation of species (Yeo et al. 2020). Recent and rapid advancements to Oxford Nanopore MinION hardware, software, and bioinformatic pipelines are quickly making this technology an affordable alternative for obtaining full-length barcodes at large-scale (Srivathsan et al. 2018, 2019, 2021). The sequencing costs for these methods are low (<0.10 USD), which is of critical importance to the democratisation of access to barcode data (Srivathsan et al. 2021). LIT can easily be implemented on datasets using fulllength, or various mini-barcodes (but see Yeo et al. 2020 for a cautionary note on the use of some mini-barcodes).

The natural next step after species delimitation will be species identification or description, depending on whether a species is new to science. LIT facilitates the description of all taxa by providing species units that have been confirmed by at least two data sources. All data acquired for LIT (barcodes, photographs, morphology, etc.) can be directly, and eventually automatically, compiled into descriptions and diagnoses. The most time-consuming step for the Swedish sample may very well be the process of determining which of the units have been described before and which are new to science. This process will be slow and laborious for regions and taxa with many described species. However, it will be much less time consuming for regions where the fauna is largely undescribed. For example, if our data were for Afrotropical phorids new species descriptions could be prepared immediately after LIT has been completed.

## Conclusion

LIT is based on a detection system that identifies where barcodes are likely to be species-specific and where they are not. This is critical not only for the future of integrative taxonomy, but also for the interpretation of metabarcoding data. We here develop and formalise an approach to validating barcode clusters as species and render the process systematic, objective, and transparent. To do so, we have developed a system to (1) identify clusters where barcodes are likely failing and (2) systematically select specimens for validation that will assure multiple species are discovered, if present. This approach integrates the power and objectivity of large-scale barcoding with targeted specialist work, thus rendering dark taxa accessible for taxonomic work. These organisms are likely critical to the functioning of our natural (and, ultimately, economic, and societal) systems; we must be committed to discovering and accurately identifying them.

---

**Supplementary Table S1.**
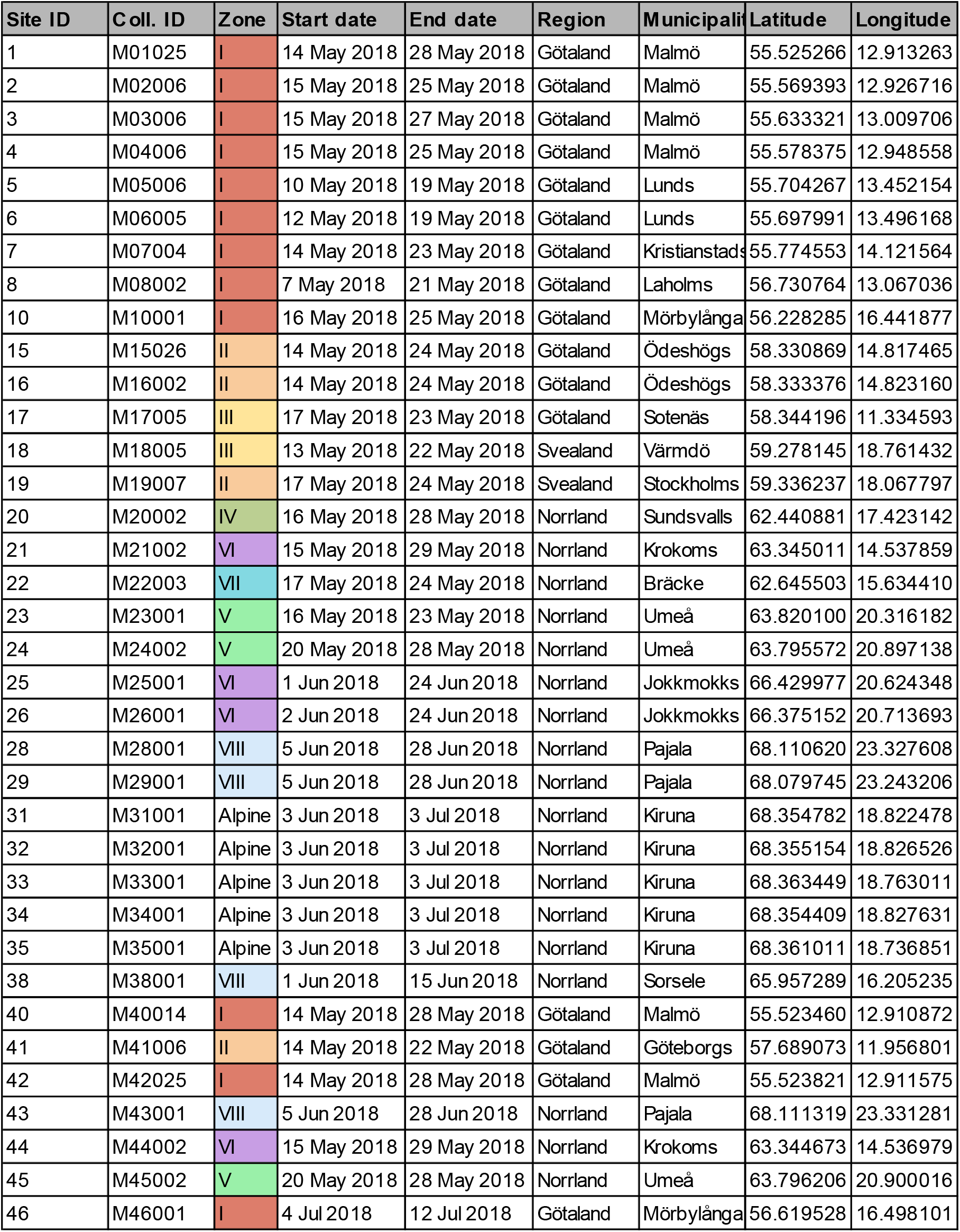
The thirty-six samples used for this study.

| Site ID | Coll. ID | Zone | Start date | End date | Region | Municipality | Latitude | Longitude |
| --- | --- | --- | --- | --- | --- | --- | --- | --- |
| 1 | M01025 | I | 14 May 2018 | 28 May 2018 | Götaland | Malmö | 55.525266 | 12.913263 |
| 2 | M02006 | I | 15 May 2018 | 25 May 2018 | Götaland | Malmö | 55.569393 | 12.926716 |
| 3 | M03006 | I | 15 May 2018 | 27 May 2018 | Götaland | Malmö | 55.633321 | 13.009706 |
| 4 | M04006 | I | 15 May 2018 | 25 May 2018 | Götaland | Malmö | 55.578375 | 12.948558 |
| 5 | M05006 | I | 10 May 2018 | 19 May 2018 | Götaland | Lunds | 55.704267 | 13.452154 |
| 6 | M06005 | I | 12 May 2018 | 19 May 2018 | Götaland | Lunds | 55.697991 | 13.496168 |
| 7 | M07004 | I | 14 May 2018 | 23 May 2018 | Götaland | Kristianstad | 55.774553 | 14.121564 |
| 8 | M08002 | I | 7 May 2018 | 21 May 2018 | Götaland | Laholms | 56.730764 | 13.067036 |
| 10 | M10001 | I | 16 May 2018 | 25 May 2018 | Götaland | Mörbylånga | 56.228285 | 16.441877 |
| 15 | M15026 | II | 14 May 2018 | 24 May 2018 | Götaland | Ödeshögs | 58.330869 | 14.817465 |
| 16 | M16002 | II | 14 May 2018 | 24 May 2018 | Götaland | Ödeshögs | 58.333376 | 14.823160 |
| 17 | M17005 | III | 17 May 2018 | 23 May 2018 | Götaland | Sotenäs | 58.344196 | 11.334593 |
| 18 | M18005 | III | 13 May 2018 | 22 May 2018 | Svealand | Värmdö | 59.278145 | 18.761432 |
| 19 | M19007 | II | 17 May 2018 | 24 May 2018 | Svealand | Stockholms | 59.336237 | 18.067797 |
| 20 | M20002 | IV | 16 May 2018 | 28 May 2018 | Norrland | Sundsvalls | 62.440881 | 17.423142 |
| 21 | M21002 | VI | 15 May 2018 | 29 May 2018 | Norrland | Krokoms | 63.345011 | 14.537859 |
| 22 | M22003 | VII | 17 May 2018 | 24 May 2018 | Norrland | Bräcke | 62.645503 | 15.634410 |
| 23 | M23001 | V | 16 May 2018 | 23 May 2018 | Norrland | Umeå | 63.820100 | 20.316182 |
| 24 | M24002 | V | 20 May 2018 | 28 May 2018 | Norrland | Umeå | 63.795572 | 20.897138 |
| 25 | M25001 | VI | 1 Jun 2018 | 24 Jun 2018 | Norrland | Jokkmokks | 66.429977 | 20.624348 |
| 26 | M26001 | VI | 2 Jun 2018 | 24 Jun 2018 | Norrland | Jokkmokks | 66.375152 | 20.713693 |
| 28 | M28001 | VIII | 5 Jun 2018 | 28 Jun 2018 | Norrland | Pajala | 68.110620 | 23.327608 |
| 29 | M29001 | VIII | 5 Jun 2018 | 28 Jun 2018 | Norrland | Pajala | 68.079745 | 23.243206 |
| 31 | M31001 | Alpine | 3 Jun 2018 | 3 Jul 2018 | Norrland | Kiruna | 68.354782 | 18.822478 |
| 32 | M32001 | Alpine | 3 Jun 2018 | 3 Jul 2018 | Norrland | Kiruna | 68.355154 | 18.826526 |
| 33 | M33001 | Alpine | 3 Jun 2018 | 3 Jul 2018 | Norrland | Kiruna | 68.363449 | 18.763011 |
| 34 | M34001 | Alpine | 3 Jun 2018 | 3 Jul 2018 | Norrland | Kiruna | 68.354409 | 18.827631 |
| 35 | M35001 | Alpine | 3 Jun 2018 | 3 Jul 2018 | Norrland | Kiruna | 68.361011 | 18.736851 |
| 38 | M38001 | VIII | 1 Jun 2018 | 15 Jun 2018 | Norrland | Sorsele | 65.957289 | 16.205235 |
| 40 | M40014 | I | 14 May 2018 | 28 May 2018 | Götaland | Malmö | 55.523460 | 12.910872 |
| 41 | M41006 | II | 14 May 2018 | 22 May 2018 | Götaland | Göteborgs | 57.689073 | 11.956801 |
| 42 | M42025 | I | 14 May 2018 | 28 May 2018 | Götaland | Malmö | 55.523821 | 12.911575 |
| 43 | M43001 | VIII | 5 Jun 2018 | 28 Jun 2018 | Norrland | Pajala | 68.111319 | 23.331281 |
| 44 | M44002 | VI | 15 May 2018 | 29 May 2018 | Norrland | Krokoms | 63.344673 | 14.536979 |
| 45 | M45002 | V | 20 May 2018 | 28 May 2018 | Norrland | Umeå | 63.796206 | 20.900016 |
| 46 | M46001 | I | 4 Jul 2018 | 12 Jul 2018 | Götaland | Mörbylånga | 56.619528 | 16.498101 |

**Supplementary Figure S1.**
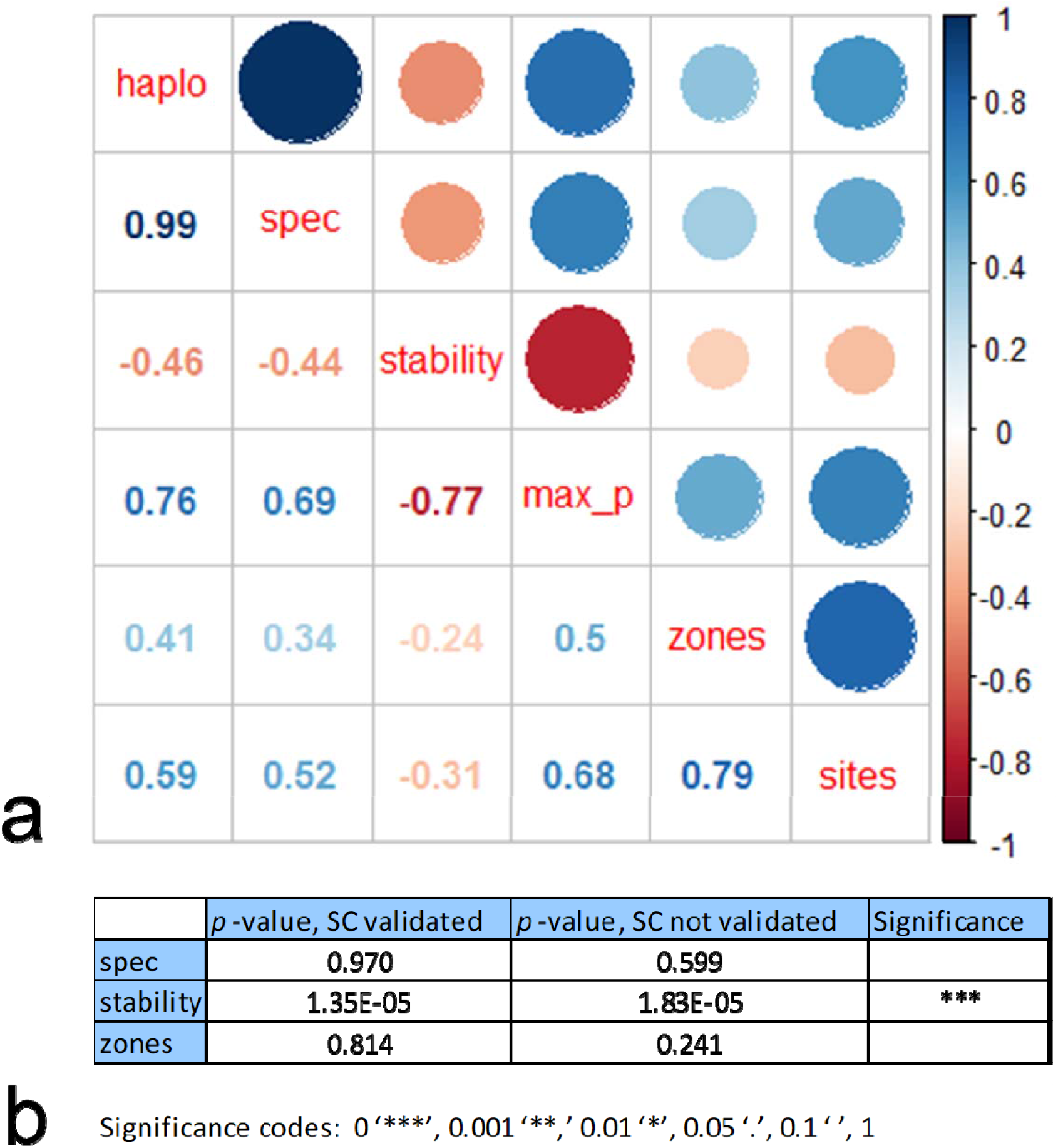
(a) Correlation matrix of explanatory variables showing strong collinearity between number of haplotypes (“haplo”) and number of specimens (“spec”), number of horticultural zones (“zones”) and number of sites (“sites”), and stability (“stability”) and maximum p-distance (“max_p”). Correlation is given numerically, and also indicated by the size and colour of the visual (see legend). (b) Significance values for our generalised linear model after collinearity was removed, showing stability as the single significant predictor of cluster failure. The model was run twice, once with species complexes (SC) counted as validated (single species) and once with SC counted as failed (multiple species).

**Supplementary Figure S2.**
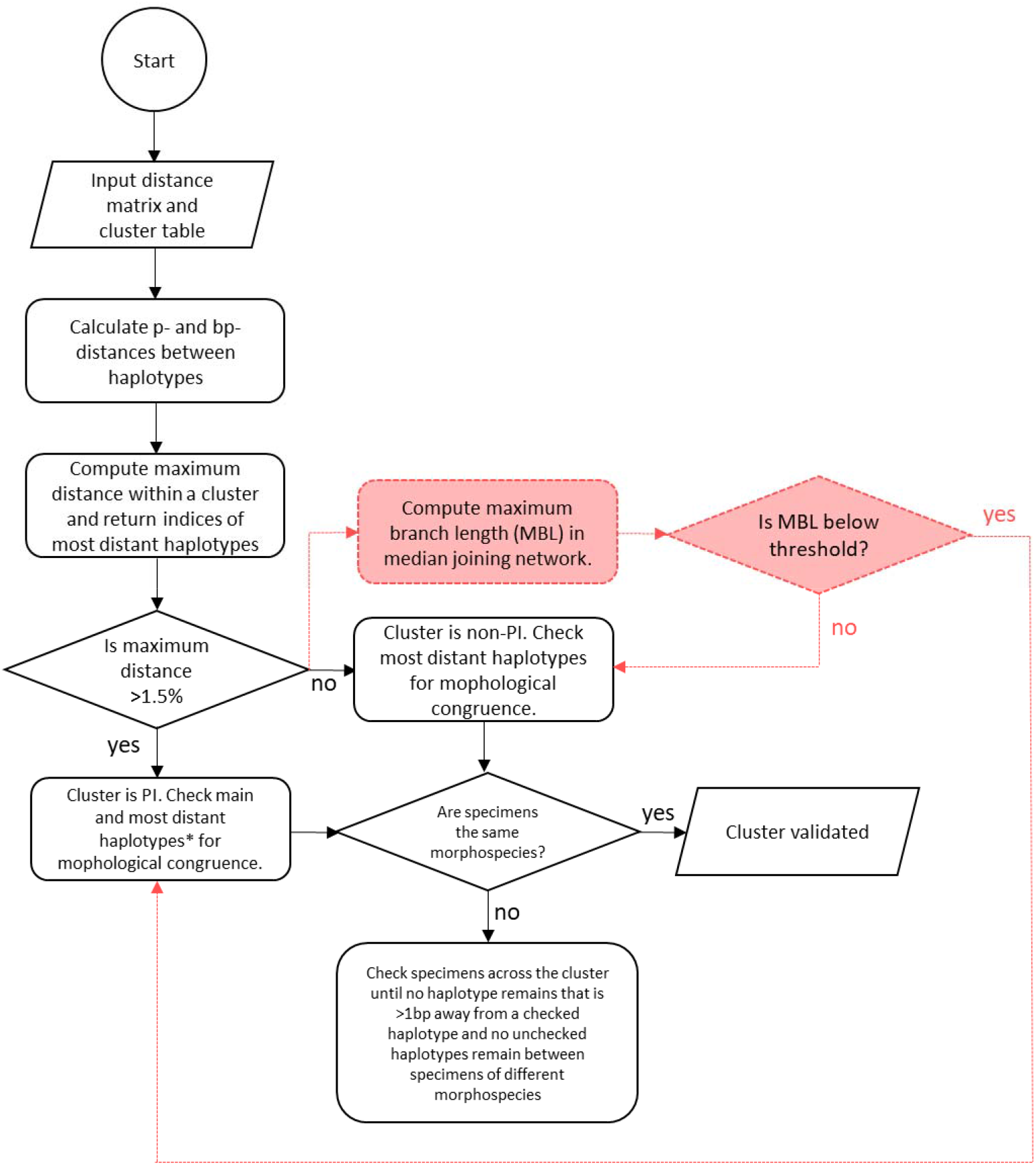
Flowchart for the specimen-picking algorithm. The algorithm was written to be applicable across clustering methods, and therefore only uses maximum p-distance to identify PI clusters. The steps in red represent a potential alternative to stability values that could be incorporated into a future version of this algorithm. *If one or both most distant haplotypes are within 2 bp of a main haplotype, the main haplotype will be checked instead

## Notes

### Competing Interest Statement

The authors have declared no competing interest.

### Summary of Updates

text changes

## References

Ahrens D., Fujisawa T., Krammer H.-J., Eberle J., Fabrizi S., Vogler A.P. 2016. Rarity and Incomplete Sampling in DNA-Based Species Delimitation. Syst. Biol. 65:17.

Andersen A., Simcox D.J., Thomas J.A., Nash D.R. 2014. Assessing reintroduction schemes by comparing genetic diversity of reintroduced and source populations: A case study of the globally threatened large blue butterfly (Maculinea arion). Biol. Conserv. 175:34–41.

Bergsten J., Bilton D.T., Fujisawa T., Elliott M., Monaghan M.T., Balke M., Hendrich L., Geijer J., Herrmann J., Foster G.N., Ribera I., Nilsson A.N., Barraclough T.G., Vogler A.P. 2012. The Effect of Geographical Scale of Sampling on DNA Barcoding. Syst. Biol. 61:851–869.

Bickel D. 2009. Why *Hilara* is not amusing: the problem of open-ended taxa and the limits of taxonomic knowledge. Diptera diversity: status, challenges, and tools. Leiden, Netherlands: E. J. Brill. p. 279–301.

Blaxter M.L. 2004. The promise of a DNA taxonomy. Philos. Trans. R. Soc. Lond. B. Biol. Sci. 359:669–679.

Brown B. 2021. Remarkably common, but undescribed, Neotropical Megaselia Rondani (Diptera: Phoridae) from Costa Rica. Submitted.

Brown B.V., Porras W. 2015. Extravagant female sexual display in a Megaselia Rondani species (Diptera: Phoridae). Biodiversity Data Journal. 3:e4368.

Butcher B.A., Smith M.A., Sharkey M.J., Quicke D.L.J. 2012. A turbo-taxonomic study of Thai Aleiodes (Aleiodes) and Aleiodes (Arcaleiodes) (Hymenoptera: Braconidae: Rogadinae) based largely on COI barcoded specimens, with rapid descriptions of 179 new species. Zootaxa. 3457:1–232.

Cesari M., Giovannini I., Bertolani R., Rebecchi L. 2011. An example of problems associated with DNA barcoding in tardigrades: a novel method for obtaining voucher specimens. Zootaxa. 3104:42.

Chapman A. 2009. Numbers of Living Species in Australia and the World (Australian Biological Resources Study, Canberra, Australia)..

Curtis T. 2006. Microbial ecologists: it’s time to “go large.” Nat. Rev. Microbiol. 4:488–488.

Dayrat B. 2005. Towards integrative taxonomy: INTEGRATIVE TAXONOMY. Biol. J. Linn. Soc. 85:407–415.

De Queiroz K. 2007. Species Concepts and Species Delimitation. Syst. Biol. 56:879–886.

Disney R.H.L. 2009. Scuttle flies (Diptera: Phoridae) Part II: the genus *Megaselia*. Fauna Arab. 24:249–357.

Eberle J., Ahrens D., Mayer C., Niehuis O., Misof B. 2020. A Plea for Standardized Nuclear Markers in Metazoan DNA Taxonomy. Trends Ecol. Evol. 35:336–345.

Geller J., Meyer C., Parker M., Hawk H. 2013. Redesign of PCR primers for mitochondrial cytochrome c oxidase subunit I for marine invertebrates and application in all-taxa biotic surveys. Mol. Ecol. Resour. 13:851–861.

Hajibabaei M., Janzen D.H., Burns J.M., Hallwachs W., Hebert P.D.N. 2006. DNA barcodes distinguish species of tropical Lepidoptera. Proc. Natl. Acad. Sci. 103:968–971.

Hartop E.A., Brown B.V. 2014. The tip of the iceberg: a distinctive new spotted-wing Megaselia species (Diptera: Phoridae) from a tropical cloud forest survey and a new, streamlined method for Megaselia descriptions. Biodivers. Data J. 2:e4093.

Hartop E.A., Brown B.V., Disney R.H.L. 2016. Flies from L.A., The Sequel: Twelve further new species of Megaselia (Diptera: Phoridae) from the BioSCAN Project in Los Angeles (California, USA). Biodivers. Data J. 4:e7756.

Hausmann A., Krogmann L., Peters R., Rduch V., Schmidt S. 2020. GBOL III: Dark Taxa. Available from https://ibol.org/barcodebulletin/research/gbol-iii-dark-taxa/.

Hebert P.D., Cywinska A., Ball S.L., deWaard J.R. 2003. Biological identifications through DNA barcodes. Proc Biol Sci. 270:313–21.

Hebert P.D.N., Braukmann T.W.A., Prosser S.W.J., Ratnasingham S., deWaard J.R., Ivanova N.V., Janzen D.H., Hallwachs W., Naik S., Sones J.E., Zakharov E.V. 2017. A Sequel to Sanger: Amplicon Sequencing That Scales..

Huemer P., Mutanen M., Sefc K.M., Hebert P.D.N. 2014. Testing DNA Barcode Performance in 1000 Species of European Lepidoptera: Large Geographic Distances Have Small Genetic Impacts. PLoS ONE. 9:e115774.

Kapli P., Lutteropp S., Zhang J., Kobert K., Pavlidis P., Stamatakis A., Flouri T. 2017. Multirate Poisson Tree Processes for single-locus species delimitation under Maximum Likelihood and Markov Chain Monte Carlo. Bioinformatics.:btx025.

Karlsson D., Hartop E.A., Forshage M., Jaschhof M., Ronquist F. 2020. The Swedish Malaise Trap Project: A 15 Year Retrospective on a Countrywide Insect Inventory. Biodivers. Data J. 8:e47255.

Katoh K., Standley D.M. 2013. MAFFT multiple sequence alignment software version 7: improvements in performance and usability. Mol. Biol. Evol. 30:772–780.

Kekkonen M., Hebert P.D.N. 2014. DNA barcode based delineation of putative species: efficient start for taxonomic workflows. Mol. Ecol. Resour. 14:706–715.

Kishan S., Marsh A. Biodiversity, Supply Chain Rank Among Biggest ESG Themes in 2021. Available from https://www.bloomberg.com/news/articles/2021-01-08/biodiversity-supply-chain-rank-among-biggest-esg-themes-in-2021.

Kumar S., Stecher G., Li M., Knyaz C., Tamura K. 2018. MEGA X: Molecular Evolutionary Genetics Analysis across Computing Platforms. Mol. Biol. Evol. 35:1547–1549.

Kwong S., Srivathsan A., Vaidya G., Meier R. 2012. Is the COI barcoding gene involved in speciation through intergenomic conflict? Mol. Phylogenet. Evol. 62:1009–1012.

Larsen B.B., Miller E.C., Rhodes M.K., Wiens J.J. 2017. Inordinate Fondness Multiplied and Redistributed: the Number of Species on Earth and the New Pie of Life. Q. Rev. Biol. 92:229–265.

Leigh J.W., Bryant D. 2015. PopART: Full-feature software for haplotype network construction. Methods Ecol Evol. 6:1110–1116.

Leray M., Yang J.Y., Meyer C.P., Mills S.C., Agudelo N., Ranwez V., Boehm J.T., Machida R.J. 2013. A new versatile primer set targeting a short fragment of the mitochondrial COI region for metabarcoding metazoan diversity: application for characterizing coral reef fish gut contents. Front. Zool. 10:34.

Locey K.J., Lennon J.T. 2016. Scaling laws predict global microbial diversity. Proc. Natl. Acad. Sci. 113:5970–5975.

Losey J.E., Vaughan M. 2006. The Economic Value of Ecological Services Provided by Insects. BioScience. 56:311.

Lücking R., Forno M.D., Moncada B., Coca L.F., Vargas-Mendoza L.Y., Aptroot A., Arias L. J., Besal B., Bungartz F., Cabrera-Amaya D.M., Cáceres M.E.S., Chaves J.L., Eliasaro S., Gutiérrez M.C., Hernández Marin J.E., de los Ángeles Herrera-Campos M., Holgado-Rojas M.E., Jonitz H., Kukwa M., Lucheta F., Madriñán S., Marcelli M.P., de Azevedo Martins S.M., Mercado-Díaz J.A., Molina J.A., Morales E.A., Nelson P.R., Nugra F., Ortega F., Paredes T., Patiño A.L., Peláez-Pulido R.N., Pérez R.E.P., Perlmutter G.B., Rivas-Plata E., Robayo J., Rodríguez C., Simijaca D.F., Soto-Medina E., Spielmann A.A., Suárez-Corredor A., Torres J.-M., Vargas C.A., Yánez-Ayabaca A., Weerakoon G., Wilk K., Pacheco M.C., Diazgranados M., Brokamp G., Borsch T., Gillevet P.M., Sikaroodi M., Lawrey J.D. 2016. Turbo-taxonomy to assemble a megadiverse lichen genus: seventy new species of Cora (Basidiomycota: Agaricales: Hygrophoraceae), honouring David Leslie Hawksworth’s seventieth birthday. Fungal Divers. 84:139–207.

Meier R., Blaimer B., Buenaventura E., Hartop E., von Rintelen T., Srivathsan A., Yeo D. 2021. A re-analysis of the data in Sharkey et al.’s (2021) minimalist revision reveals that BINs do not deserve names, but BOLD Systems needs a stronger commitment to open science. BioRxiv Prepr.

Meier R., Shiyang K., Vaidya G., Ng P.K. 2006. DNA barcoding and taxonomy in Diptera: a tale of high intraspecific variability and low identification success. Syst Biol. 55:715–28.

Meier R., Wong W., Srivathsan A., Foo M. 2016. $1 DNA barcodes for reconstructing complex phenomes and finding rare species in specimen-rich samples. Cladistics. 32:100–110.

Meier R., Zhang G., Ali F. 2008. The Use of Mean Instead of Smallest Interspecific Distances Exaggerates the Size of the “Barcoding Gap” and Leads to Misidentification. Syst. Biol. 57:809–813.

Mora C., Tittensor D.P., Adl S., Simpson A.G., Worm B. 2011. How many species are there on Earth and in the ocean? PLoS Biol. 9:e1001127.

Padial J.M., Miralles A. 2010. The integrative future of taxonomy.:14.

Page R. 2011. Dark taxa: GenBank in a post-taxonomic world. Available from https://iphylo.blogspot.com/2011/04/dark-taxa-genbank-in-post-taxonomic.html.

Page R.D. 2016. DNA barcoding and taxonomy: dark taxa and dark texts. Philos Trans R Soc Lond B Biol Sci. 371.

Pante E., Schoelinck C., Puillandre N. 2015. From Integrative Taxonomy to Species Description: One Step Beyond. Syst. Biol. 64:152–160.

Pentinsaari M., Salmela H., Mutanen M., Roslin T. 2016. Molecular evolution of a widely-adopted taxonomic marker (COI) across the animal tree of life. Sci. Rep. 6:35275.

Pérez-Silva J.G., Araujo-Voces M., Quesada V. 2018. nVenn: generalized, quasi-proportional Venn and Euler diagrams. Bioinformatics. 34:2322–2324.

Puillandre N., Lambert A., Brouillet S., Achaz G. 2012. ABGD, Automatic Barcode Gap Discovery for primary species delimitation: ABGD, AUTOMATIC BARCODE GAP DISCOVERY. Mol. Ecol. 21:1864–1877.

Puillandre N., Modica M.V., Zhang Y., Sirovich L., Boisselier M.-C., Cruaud C., Holford M., Samadi S. 2012. Large-scale species delimitation method for hyperdiverse groups: LARGE-SCALE SPECIES DELIMITATION. Mol. Ecol. 21:2671–2691.

Ratnasingham S., Hebert P.D. 2013. A DNA-based registry for all animal species: the barcode index number (BIN) system. PLoS One. 8:e66213.

Riedel A., Sagata K., Suhardjono Y.R., Tänzler R., Balke M. 2013. Integrative taxonomy on the fast track - towards more sustainability in biodiversity research. Front. Zool. 10:1–9.

Riksförbundet Svensk Trädgård. 2018. Zonkartan. Available from http://www.tradgard.org/svensk_tradgard/zonkartan.html.

Schlick-Steiner B.C., Steiner F.M., Seifert B., Stauffer C., Christian E., Crozier R.H. 2010. Integrative Taxonomy: A Multisource Approach to Exploring Biodiversity. Annu. Rev. Entomol. 55:421–438.

Sites J.W., Marshall J.C. 2004. Operational Criteria for Delimiting Species. Annu. Rev. Ecol. Evol. Syst. 35:199–227.

Smith M.A., Rodriguez J.J., Whitfield J.B., Deans A.R., Janzen D.H., Hallwachs W., Hebert P.D.N. 2008. Extreme diversity of tropical parasitoid wasps exposed by iterative integration of natural history, DNA barcoding, morphology, and collections. Proc. Natl. Acad. Sci. 105:12359–12364.

Soviċ I., Šikić M., Wilm, A. Fenlon, S.N., Chen, S., Nagarajan, N. (2016). Fast and sensitive mapping of nanopore sequencing reads with GraphMap. Nat. Comm. 7:11307.

Srivathsan A., Baloğlu B., Wang W., Tan W.X., Bertrand D., Ng A.H.Q., Boey E.J.H., Koh J.J.Y., Nagarajan N., Meier R. 2018. A MinION™-based pipeline for fast and costeffective DNA barcoding. Mol. Ecol. Resour. 18:1035–1049.

Srivathsan A., Hartop E.A., Puniamoorthy J., Lee W.T., Kutty S.N., Kurina O., Meier R. 2019. Rapid, large-scale species discovery in hyperdiverse taxa using 1D MinION sequencing. BMC Biol. 17:96.

Srivathsan A., Lee L., Katoh K., Hartop E., Narayanan Kutty S., Wong J., Yeo D., Meier R. 2021. MinION barcodes: biodiversity discovery and identification by everyone, for everyone. BioRxiv Prepr.

Stamatakis A. 2014. RAxML Version 8: A tool for Phylogenetic Analysis and Post-Analysis of Large Phylogenies. Bioinformatics.

Swiss Re Institute. 2020. Biodiversity and Ecosystem Services: A business case for re/insurance..

Tautz D., Arctander P., Minelli A., Thomas R.H., Vogler A.P. 2003. A plea for DNA taxonomy. Trends Ecol. Evol. 18:70–74.

Thomas J.A. 1995. The ecology and conservation of Maculinea arion and other European species of large blue butterfly. In: Pullin A.S., editor. Ecology and Conservation of Butterflies. Dordrecht: Springer Netherlands. p. 180–197.

Townes H. 1972. A light-weight Malaise trap. Entomol. News. 83:239–247.

Truett G.E., Heeger P., Mynatt R.L., Truett A.A., Walker J.A., Warman M.L. 2000. Preparation of PCR-Quality Mouse Genomic DNA with Hot Sodium Hydroxide and Tris (HotSHOT). BioTechniques. 29:52–54.

Vaser R., Sović I., Nagarajan N., Šikić M. 2017. Fast and accurate de novo genome assembly from long uncorrected reads. Genome Res. 27:737–746.

Vitecek S., Kučinić M., Previšiċ A., Živiċ I., Stojanoviċ K., Keresztes L., Bálint M., Hoppeler F., Waringer J., Graf W., Pauls S.U. 2017. Integrative taxonomy by molecular species delimitation: multi-locus data corroborate a new species of Balkan Drusinae micro-endemics. BMC Evol. Biol. 17:129.

Vogler A.P., Monaghan M.T. 2007. Recent advances in DNA taxonomy. J. Zool. Syst. Evol. Res. 45:1–10.

Wang W.Y., Srivathsan A., Foo M., Yamane S., Meier R. 2018. Sorting specimen-rich invertebrate samples with cost-effective NGS barcodes: validating a reverse workflow for specimen processing. Mol Ecol Resour.

Wiens J.J. 2007. Species Delimitation: New Approaches for Discovering Diversity. Syst. Biol. 56:875–878.

Will K.W., Mishler B.D., Wheeler Q.D. 2005. The Perils of DNA Barcoding and the Need for Integrative Taxonomy. Syst. Biol. 54:844–851.

Yeo D., Srivathsan A., Meier R. 2020. Longer is not always better: Optimizing barcode length for large-scale species discovery and identification. Syst. Biol.:syaa014.

Yong E. 2009 How research save the Large Blue butterfly. Available from https://www.nationalgeographic.com/science/article/how-research-saved-the-large-blue-butterfly.

Zhang J., Kapli P., Pavlidis P., Stamatakis A. 2013. A general species delimitation method with applications to phylogenetic placements. Bioinformatics. 29:2869–2876.

Zhang J., Kobert K., Flouri T., Stamatakis A. 2014. PEAR: a fast and accurate Illumina Paired-End reAd mergeR. Bioinformatics. 30:614–620.

